# Gain-of-function mutation in *SKAP2* leads to type 1 diabetes and broader autoimmunity through hyperactive integrin signaling in myeloid cells

**DOI:** 10.64898/2026.04.02.716136

**Authors:** Courtney M. Tamaki, Chester E. Chamberlain, Clare L. Abram, Sumith Poojary, Jennifer Bridge, Jennifer L. Matsuda, Whitney Tamaki, Niklas Rutsch, Lauren Spector, Wesley Dixon, Irina Proekt, Lisa R. Letourneau-Freiberg, Louis H. Philipson, Michael S. German, Mark S. Anderson, Clifford A. Lowell

## Abstract

Many genetic variants associated with increased type 1 diabetes (T1D) risk are located within the *SKAP2* gene; however, the mechanisms by which these variants confer disease risk remain unclear. *SKAP2* encodes an adapter protein that functions within the integrin signaling pathway and is found at the highest levels in myeloid leukocytes. We recently identified a *de novo* gain-of-function *SKAP2* mutation in an individual with T1D, leading to hyperactive integrin signaling in myeloid cells. To dissect the mechanisms by which this mutation may lead to T1D, we generated a knock-in mouse line containing the orthologous p.G153R substitution in mouse SKAP2 on the diabetes-prone nonobese diabetic (NOD) genetic background. Both female and male SKAP2^G153R/G153R^ mice developed accelerated T1D. The SKAP2^G153R/G153R^ mice also exhibited a unique spectrum of autoantibodies, leading to immune-complex nephritis. Accelerated infiltration of pancreatic islets by myeloid cells, B lymphocytes, and activated T cells was observed in SKAP2^G153R/G153R^ mice. Single-cell RNA sequencing demonstrated a type 1 IFNγ-driven inflammatory program within the pancreatic islets of SKAP2^G153R/G153R^ mice. Dendritic cells from SKAP2^G153R/G153R^ mice demonstrated increased antigen-presenting capacity, characterized by enhanced adhesion to T cells during immune synapse formation. Macrophages and neutrophils from SKAP2^G153R/G153R^ mice also showed increased integrin signaling responses, with neutrophils expressing high levels of activated β2 integrins on the cell surface. When backcrossed onto the C57BL/6J genetic background, the SKAP2^G153R/G153R^ mice developed spontaneous autoantibody formation and exhibited accelerated autoimmunity, including nephritis, in the pristane-induced model of autoimmune disease. These findings demonstrate that dysregulation of leukocyte integrin signaling, through alterations in *SKAP2,* may increase the genetic risk for autoimmunity and T1D.

## INTRODUCTION

Type 1 diabetes (T1D) is a complex autoimmune disease characterized by T cell-mediated destruction of pancreatic β cells, leading to insulin deficiency. Many individuals with T1D often manifest other autoimmune processes, such as Hashimoto’s thyroiditis, Addison’s disease, and Sjögren’s syndrome, frequently accompanied by disease-defining autoantibodies to specific tissue antigens (Frommer and Kahaly, 2020). According to family studies, T1D has an approximately 50% hereditary risk (Noble and Erlich, 2012). Most of this genetic susceptibility has been linked to genes in the major histocompatibility complex (HLA). Still, many genome-wide association studies (GWAS) have linked T1D to other, often immune-regulatory, gene variants (Minniakhmetov et al., 2024; Noble and Erlich, 2012). Examples of such immune-regulatory genes include *CTLA-4*, *TNFAIP3*, *STAT3*, *IL-6R*, and others (Ferreira et al., 2013; Fung et al., 2009; Velayos et al., 2017; Wang et al., 2014). One of the most common gene variants associated with T1D is within the *SKAP2* locus (Haris et al., 2021; Krischer et al., 2022; Reddy et al., 2011; Robertson et al., 2021). In perhaps the most extensive genetic study of T1D patients to date (61,427 participants), the *SKAP2* locus was identified as having the second largest number of single-nucleotide polymorphisms (SNPs), just behind *IL-27*, estimated to be present in nearly 30% of T1D patients (Robertson et al., 2021). *SKAP2* variants were also the second-most-common gene SNPs identified in a Qatari cohort (1096 patients), accounting for 5% of all individuals (Haris et al., 2021). Variants in *SKAP2* are also associated with other autoimmune diseases, including Crohn’s disease (Garza-Hernandez et al., 2022; Liu et al., 2015) and inflammatory venous insufficiency (Ellinghaus et al., 2017). The molecular mechanisms by which these *SKAP2* SNPs contribute to T1D susceptibility or broader autoimmunity remain unclear.

Src-kinase-associated protein 2 (SKAP2 – previously known as SKAP-HOM) is an intracellular signaling scaffolding protein that links extracellular receptors, particularly integrin adhesion receptors, to the actin cytoskeleton (reviewed in Wilmink and Spalinger, 2023). SKAP2 is found primarily in myeloid leukocytes, at lower levels in B lymphocytes, and at low levels in neurons, some fibroblast types, and other stromal cells (Immgen.org). The homolog of SKAP2, i.e., SKAP1, is expressed primarily in T lymphocytes, natural killer (NK) cells, and innate lymphoid cells (ILCs), where it regulates integrin activation, cell migration, and optimizes T cell proliferation (reviewed in Liu et al., 2023). Similarly, SKAP2 plays a central role in regulating myeloid cell recruitment and extravasation into inflammatory sites by linking actin regulatory proteins, such as the Wiskott–Aldrich syndrome protein (WASP) complex, to the plasma membrane while also directly associating with integrin-binding molecules (such as Talin-1) to facilitate integrin signaling. Macrophages from SKAP2-deficient mice have impaired adhesion to extracellular matrix proteins (such as fibronectin), impaired actin remodeling, and reduced chemotaxis responses (Alenghat et al., 2012). Neutrophils from *Skap2^-/-^* mice exhibit impaired integrin β2 activation, resulting in inadequate adhesion-dependent stimulation of the oxidative burst and reduced recruitment to inflammatory sites (Boras et al., 2017). As a result, *Skap2^-/-^* mice have increased susceptibility to bacterial and fungal infections (Nguyen et al., 2020; Nguyen et al., 2021). Similar results have recently been demonstrated in neutrophil-like cell lines, showing that SKAP2 deficiency leads to a variety of functional defects in neutrophils downstream of the CD11b/CD18 integrin receptors (Bouti et al., 2024). Dendritic cells from *Skap2^-/-^* mice exhibit reduced adhesion to T cells, leading to poor priming of antigen-specific T cells. This has been described as a mechanism to explain the reduced inflammatory responses observed in *Skap2^-/-^* mice in the experimental autoimmune encephalitis model (Reinhold et al., 2009; Togni et al., 2005). Reduced levels of SKAP2 have been observed in acute myeloid leukemia lines, which correlated with reduced production of radical oxygen species (Hu et al., 2025).

The SKAP2 protein has three well-described domains: an N-terminal dimerization domain (DM), followed by a pleckstrin homology (PH) domain, which is required for the association of SKAP2 with the plasma membrane, and a C-terminal SH3 domain, which is necessary for association with signaling molecules such as WASP and others (Levillayer et al., 2023). SKAP2 is held in an auto-inhibited state in resting cells through intramolecular binding of the DM and PH domains. Cell activation leads to tyrosine phosphorylation of SKAP2, resulting in dissociation of the DM and PH domains. This allows the PH domain to bind to plasma membrane phosphatidylinositol-3,4,5-triphosphate (PIP3) lipids, while also facilitating the association of the SH3 domain with WASP and other proteins (Swanson et al., 2008). Interestingly, disrupting the DM-PH interaction through mutation (D129K substitution within the PH domain) leads to unregulated binding to PIP3 lipids, resulting in hyperactive actin polymerization and increased adhesion in macrophages (Swanson et al., 2008).

We previously identified a 24-year-old woman with T1D associated with autoimmune hypothyroidism, scleroderma, and multiple associated autoantibodies (Rutsch et al., 2021). Whole-exome sequencing of the woman and her direct relatives revealed a *de novo* heterozygous mutation, resulting in a single amino acid substitution (p.G153R) in the PH domain of SKAP2 (SKAP2 G153R). This mutation resulted in hyperactive downstream integrin signaling in peripheral blood myeloid leukocytes. Indeed, resting monocyte-derived macrophages from the individual showed increased association of SKAP2 and WASP with the plasma membrane and enhanced downstream integrin signaling. These observations suggested that the G153R mutation produced a gain-of-function, constitutively active form of SKAP2, which may drive autoimmunity. To test this, we generated a knock-in mouse model on the nonobese diabetic (NOD) genetic background carrying the orthologous SKAP2 G153R mutation. These knock-in mice exhibited accelerated T1D, characterized by rapid insulitis associated with broad-spectrum autoimmunity. We used these mice to define the mechanisms by which hyperactive integrin signaling in myeloid leukocytes may drive autoimmune T1D. Our observations suggest that this *SKAP2* gain-of-function mutation leads to enhanced myeloid cell activation and dysregulated antigen presentation, which facilitates the development of autoimmune disease.

## RESULTS

### Generation and validation of SKAP2^G153R^ knock-in allele in NOD mice

We used CRISPR/Cas9 mutagenesis to introduce a single point mutation into the mouse *Skap2* gene that results in the substitution of glycine 153 to arginine (p.G153R – referred to as SKAP2^G153R^ knock-in allele throughout) (Fig. 1A). The mutation was introduced directly into mouse embryos from the nonobese diabetic (NOD) strain, a well-established animal model of spontaneous autoimmune diabetes which has been widely used to study defects in immune tolerance (Anderson and Bluestone, 2005; Khosravi-Maharlooei et al., 2022; Warshauer et al., 2021). All mice used in this study were maintained on the NOD genetic background, unless otherwise specified. Western blot analysis confirmed SKAP2 expression in murine monocytes, dendritic cells, and neutrophils, with little to no expression detected in lymphoid B or T cells. The SKAP2 G153R protein did not have an altered cellular expression or abundance compared to WT SKAP2 (Fig. 1B), as observed for the human ortholog (Rutsch et al., 2021). Since SKAP2 is tyrosine phosphorylated in response to integrin engagement (Boras et al., 2017; Timms et al., 1999), we performed immunoprecipitation experiments to assess the tyrosine phosphorylation level of SKAP2 G153R. While overall protein tyrosine phosphorylation levels in whole bone marrow cells were similar between SKAP2^+/+^ and SKAP2^G153R/G153R^ knock-in mice, SKAP2 G153R exhibited increased tyrosine phosphorylation compared to WT SKAP2 (Fig. 1C), suggesting that the mutation leads to an activated form of the protein.

**Figure 1.**
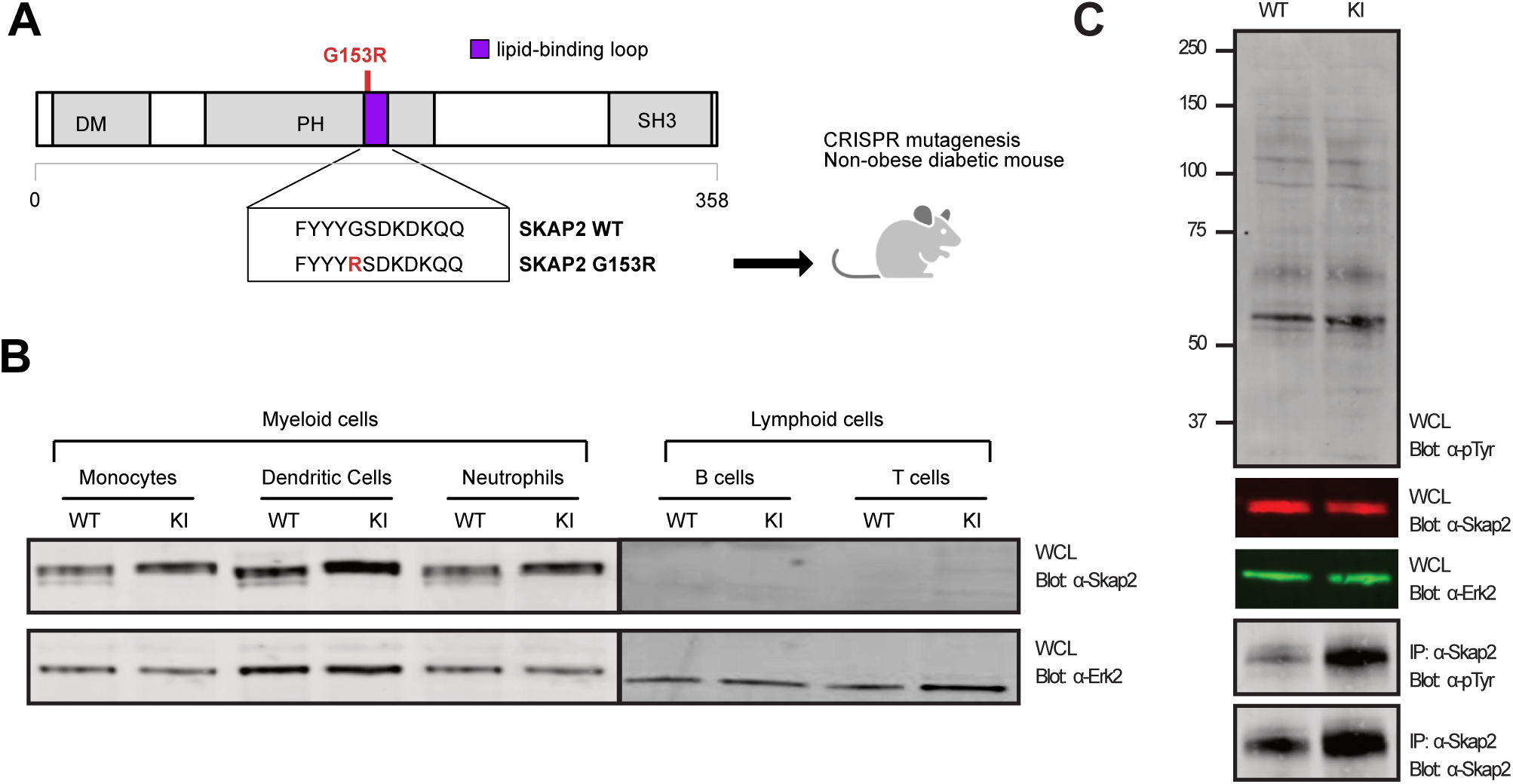
Generation of SKAP2^G153R^ knock-in allele in NOD mice. **(A)** SKAP2 G153R mutation located in the lipid-binding loop of the PH domain was introduced into the nonobese diabetic (NOD) mouse line using CRISPR/Cas9. **(B)** Western blot showing SKAP2 protein levels in whole cell lysates of the sorted cell lineages CD11b^+^ monocytes, CD11c+ dendritic cells, Ly6G^+^ neutrophils, CD19^+^ B cells, and CD3^+^ T cells from NOD female mice that were SKAP2*^+/+^* (WT) or SKAP*2*^G153R/G153R^ (labeled as KI) immunoblotted with anti-SKAP2 (top) and anti-ERK2 (bottom) as a control. **(C)** Whole bone marrow cell lysates from WT or KI mice were immunoblotted directly with anti-phospho-Tyr (labeled PTyr), anti-SKAP2, or anti-ERK2. Lysates were subjected to immunoprecipitation (IP) using anti-SKAP2, followed by immunoblotting with anti-PTyr or anti-SKAP2. Results are representative of three independent experiments.

### The SKAP2^G153R^ knock-in allele accelerates autoimmune diabetes and broad autoimmunity

To evaluate whether the SKAP2^G153R^ knock-in allele promotes T1D, blood glucose levels were measured weekly for 30-40 weeks in cohorts of NOD mice that were either wild-type (SKAP2*^+/+^*or WT), heterozygous (SKAP2^G153R/+^), or homozygous (SKAP2^G153R/G153R^) for the SKAP2^G153R^ knock-in allele. Male and female mice were analyzed separately because of known sex-dependent differences in diabetes onset in the NOD model (Anderson and Bluestone, 2005). Diabetes onset was defined as blood glucose concentrations exceeding 200 mg/dl on two successive measurements obtained one week apart. SKAP2^G153R/G153R^ female mice exhibited a marked acceleration in diabetes onset, with 50% of the animals becoming diabetic by 15 weeks of age compared to 20 weeks in WT females (Fig. 2A). In contrast, heterozygous SKAP2^G153R/+^ females developed diabetes at the same rate as WT controls (Fig. S1A). Among males, both homozygous and heterozygous mice developed diabetes with an incidence of ∼50% at 30 weeks of age (Fig. 2B and S1B), while none of the WT males developed diabetes. RAG2-deficient SKAP2^G153R/G153R^ mice, which lack mature B and T cells, were completely protected from diabetes, demonstrating that adaptive immunity was required for diabetes development (Fig. 2A, B) (Warshauer et al., 2021). Given the minimal expression of SKAP2 in lymphocytes, this suggests the SKAP2^G153R^ knock-in allele likely affects other immune cells, such as antigen-presenting myeloid leukocytes, to enhance the pathogenic adaptive immune responses that drive pancreatic islet cell destruction.

**Figure 2.**
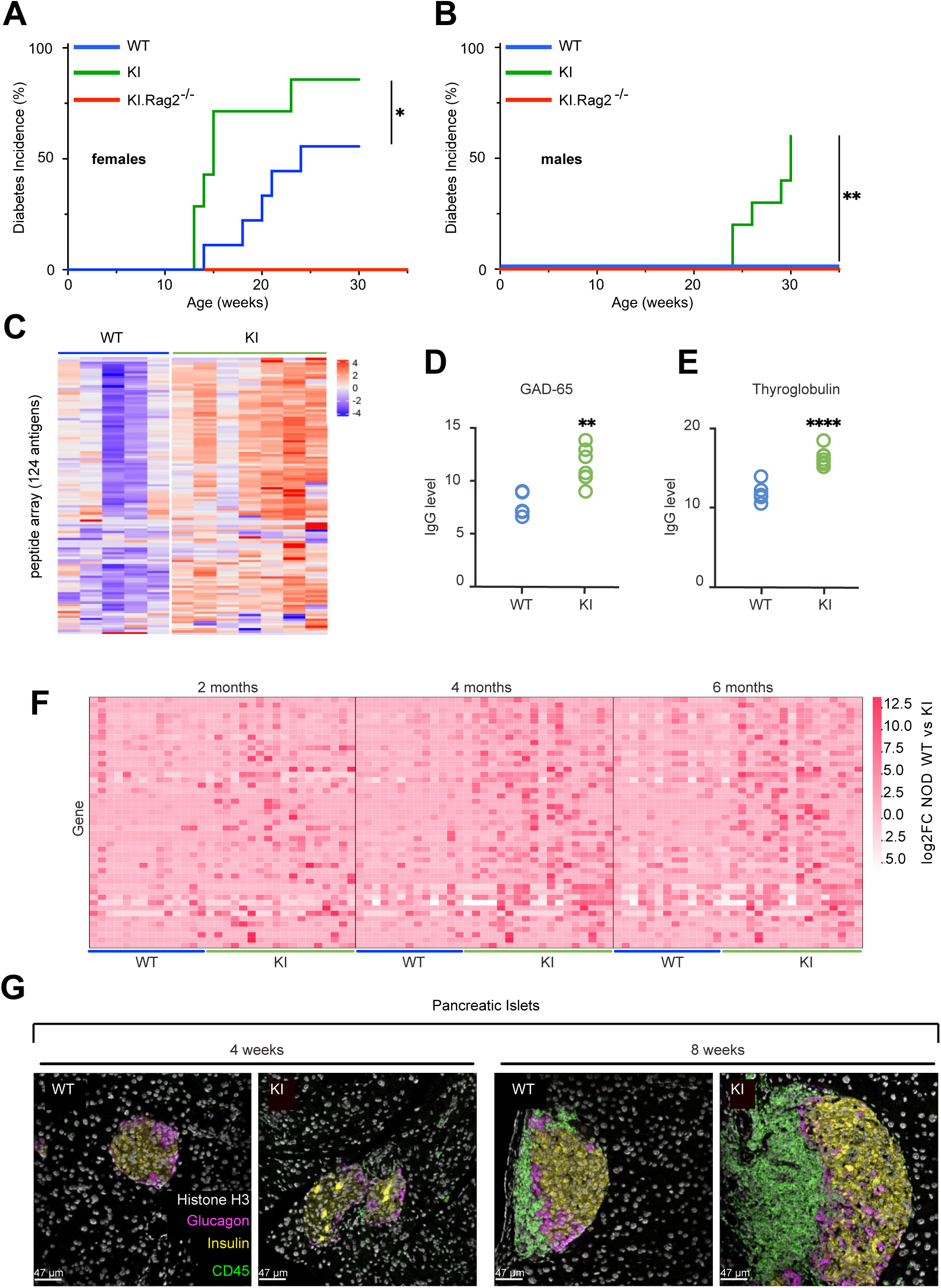
The SKAP2 G153R mutation accelerates autoimmune diabetes. **(A,B)** Cohorts of NOD mice that were SKAP2^+/+^ (WT), SKAP2^G153R/G153R^ (labeled as KI) or Rag2-deficient SKAP2^G153R/G153R^ (KI.Rag2^-/-^) were monitored for blood glucose levels weekly. Diabetes (blood glucose > 200 mg/dl on two tests separated by a week) onset and incidence over 30 – 40 weeks are shown in female **(A)** and male **(B)** mice (females: WT, n=9; KI, n=10; KI.Rag2^-/*-*^, n=5; males: WT, n=10; KI, n=10; KI.Rag2^-/*-*^, n=5). **(C-E)** Serum samples from twelve-week-old female WT (n=5) or SKAP2^G153R/G153R^ (KI) (n=7) mice were analyzed for IgG autoantibodies by tissue antigen array (see Table S1 for a list of antigens displayed). Data are plotted as a heat map relative to IgG control, with antibody score defined as log_2_ fold signal/signal-to-noise+1. Specific values for GAD-65 and thyroglobulin in WT versus KI samples are shown. **(F)** Serum samples from female WT (n=13) or SKAP2^G153R/G153R^ (KI) (n=18) mice at ages two, four and six months were analyzed for autoantibodies by PhIP-Seq. Heatmaps showing gene-level log₂ fold-change (log₂FC) values between WT versus KI. Only genes exhibiting a log₂FC > 1 at any time point are shown (see Table S2 for a list of antigens displayed). For genes represented by multiple peptides, log₂FC values were averaged to derive a single gene-level enrichment score. **(G)** Pancreatic islets from four and eight-week-old female WT or SKAP2^G153R/G153R^ (KI) mice were stained with a metal-conjugated 22 antibody panel (see Materials and Methods for a list of antibodies used for MIBI imaging). Scales are 47um. The markers shown demonstrate the difference in CD45^+^ (green) immune cell infiltration, pancreatic insulin (yellow)- or glucagon (magenta)-secreting islet cells, or surrounding exocrine pancreatic cells, as indicated by histone H3 (white) staining. Diabetes onset curves were analyzed using Log-rank testing (Mantel-Cox test); *=p<0.05; **=p<0.01; ****=p<0.0001.

To further assess whether the SKAP2 G153R gain-of-function mutation enhances pathogenic T cell responses, we crossed the SKAP2^G153R/G153R^ mice to BDC2.5 T cell receptor transgenic mice, which express a CD4^+^ TCR specific for a chromogranin A-derived peptide expressed by pancreatic islet cells (Stadinski et al., 2010). Despite the presence of a highly diabetogenic TCR, BDC2.5 transgenic mice rarely exhibit diabetes due to the development of regulatory CD4^+^ cells (T_reg_) that restore peripheral immune tolerance (Lepault and Gagnerault, 2000; Stadinski et al., 2010; Tang et al., 2004). Strikingly, heterozygous SKAP2^G153R/+^ females carrying the BDC2.5 transgene developed diabetes at approximately the same rate as non-transgenic homozygous SKAP2^G153R/G153R^ animals (Fig. S1C). This result indicates that a single copy of the SKAP2^G153R^ allele is sufficient to overcome T_reg_-mediated suppression of chromogranin A-specific T effector cells, thereby promoting the development of autoimmune diabetes.

To evaluate the extent of autoimmunity in SKAP2^G153R/G153R^ mice, we examined serum autoantibody levels in female mice using a multiplex protein array. At twelve weeks of age, SKAP2^G153R/G153R^ mice exhibited elevated autoantibody reactivity compared to WT mice across a broad spectrum of self-antigens (Fig. 2C and Table S1). Notably, this included increased titers of islet-specific glutamic acid decarboxylase 65 (GAD-65) autoantibodies and anti-thyroglobulin antibodies (Fig. 2D, E), similar to the autoantibody profile observed in the patient harboring the *SKAP2* mutation (Rutsch et al., 2021). Twelve-week-old male SKAP2^G153R/G153R^ knock-in mice also showed elevated autoantibodies compared to WT controls, though the magnitude was reduced compared with that observed in females (not shown).

To validate and extend the protein array findings, we also used a recently developed murine proteome-wide phage display library platform (PhiP-seq) to identify autoantibody specificities (Rackaityte et al., 2023). This highly sensitive method has been widely used to profile autoantibody repertoires in human autoimmune diseases (Mandel-Brehm et al., 2022; Vazquez et al., 2020). Using this approach, we confirmed that SKAP2^G153R/G153R^ female mice displayed a wide range of autoantibody reactivity to various peptides encoded by a diverse range of genes, as early as 2 months of life, in contrast to age-matched WT female mice (Fig. 2F and Table S2). Although the autoantibody specificities identified by PhiP-seq differed from those detected by protein array analysis due to fundamental differences in peptide presentation between the two platforms, both approaches consistently demonstrate that the SKAP2 G153R mutation drives the early development of broad-spectrum autoimmunity.

The development of pancreatic insulitis in SKAP2^G153R/G153R^ versus WT female mice was assessed by immunohistochemical methods, using both light microscopy and multiplexed ion beam imaging (MIBI). Pancreatic sections from four to twelve-week-old mice were stained with antibodies to detect T cells, B cells, macrophages, and dendritic cells. At all early time points examined, SKAP2^G153R/G153R^ mice exhibited more extensive immune cell infiltration of pancreatic islets compared to WT animals (Fig. S2A, B). Quantitative analysis revealed significantly higher numbers of dendritic cells and macrophages in islets at six or eight weeks of age. In contrast, B cell infiltration was more pronounced at eight to twelve weeks. By twenty weeks of age, both the SKAP2^G153R/G153R^ and WT mice displayed near complete immune-mediated destruction of the pancreatic islets and were indistinguishable (not shown). Accelerated insulitis in SKAP2^G153R/G153R^ mice was independently confirmed by MIBI microscopy, staining for CD45^+^ immune cells, as well as insulin- and glucagon-producing cells. At eight weeks of age, pancreatic islets from SKAP2^G153R/G153R^ mice had a strikingly increased number of CD45^+^ cells compared to WT controls (Fig. 2G). Together, these data demonstrate that the SKAP2^G153R/G153R^ gain-of-function mutation accelerates immune cell infiltration and destruction of pancreatic islets, providing a basis for the earlier onset of diabetes observed in these animals.

In addition to pancreatic autoimmunity, NOD mice are predisposed to inflammation in other tissues, including the thyroid and salivary glands (Aubin et al., 2022). Despite this, immune complex glomerulonephritis is not a significant feature in NOD mice. To determine whether the expanded autoantibody repertoire observed in female SKAP2^G153R/G153R^ mice resulted in immune complex deposition in the kidney, renal sections from seventeen-week-old female WT and SKAP2^G153R/G153R^ mice were stained for IgG, IgM, and complement C3. Female SKAP2^G153R/G153R^ mice exhibited substantial immune complex deposition within the kidney glomeruli compared to WT animals, accompanied by inflammatory cell infiltration, as revealed by H&E histology (Fig. S2C, D). In addition, SKAP2^G153R/G153R^ mice showed a trend towards increased salivary gland inflammation, although the severity varied between animals (Fig. S2E, F). No significant inflammatory infiltrates were detected in the lung, liver, or intestine of either WT or SKAP2^G153R/G153R^ mice. These findings suggest that the SKAP2^G153R/G153R^ mutation exacerbates extra-pancreatic inflammatory manifestations of the NOD mouse model.

To define early systemic and tissue-specific immune alterations associated with the SKAP2 G153R gain-of-function mutation, peripheral lymphoid tissues and pancreatic islets were analyzed by spectral flow cytometry using a 28-marker antibody panel (Table S4). Female SKAP2^G153R/G153R^ and WT mice at four and eight weeks of age were used to capture immune cell changes before the onset of frank diabetes. Immunophenotyping CD45^+^ leukocytes from bone marrow, spleen, and pancreatic lymph nodes revealed broadly comparable distributions of myeloid and lymphoid cell populations between WT and SKAP2^G153R/G153R^ mice at both time points. The only significant difference detected was an increased frequency of F4/80^+^ macrophages in the bone marrow of four-week-old SKAP2^G153R/G153R^ animals (Fig. S3A). In contrast, more pronounced differences were evident within the islets. At four weeks of age, islet-infiltrating macrophages from SKAP2^G153R/G153R^ mice exhibited a lower expression level of CD206, a marker of M2 macrophages (Nevarez-Mejia et al., 2024), and a higher percentage of CD4^+^ effector T cells (defined as CD44^+^CD62L^-^) with a concomitant reduction of CD69^+^ cells (Fig. S3C, D). Markers of dendritic cell activation were broadly similar between WT versus SKAP2^G153R/G153R^ animals in both islets and pancreatic draining lymph nodes. However, conventional dendritic cells (cDCs) isolated from SKAP2^G153R/G153R^ islets showed elevated PD-L1 expression compared to WT controls (Fig. S3E, F). Collectively, these findings indicate that early inflammatory changes in SKAP2^G153R/G153R^ mice are predominantly localized to the pancreatic islets and characterized by increased numbers of activated leukocytes at this site rather than by widespread systemic immune dysregulation. The elevated expression of PD-L1 on islet cDCs in four-week-old SKAP2^G153R/G153R^ mice is consistent with early exposure to IFNγ, which is known to drive up PD-L1 expression on cDCs (Peng et al., 2020).

### The SKAP2 G153R mutation triggers early IFNγ-driven inflammatory response in islet myeloid cells

To obtain a high-resolution view of pancreatic insulitis, we performed single-cell RNA sequencing (scRNAseq) of CD45^+^ immune cells isolated from pancreatic islets from four and eight-week-old WT and SKAP2^G153R/G153R^ female mice. Approximately 20,000 CD45^+^ cells were profiled per condition, except for four-week-old WT mice, from which only 3,000 cells could be recovered due to the low abundance of islet-infiltrating immune cells at this early time point. Unsupervised clustering and visualization were performed using Seurat (Fig. 3A) (Stuart et al., 2019). Seventeen distinct immune populations were identified based on canonical marker gene, consistent with previous single-cell analyses of NOD islet inflammation (Fig. S4A) (Zakharov et al., 2020). At four weeks of age, islets from SKAP2^G153R/G153R^ mice had a marked expansion of infiltrating CD45^+^ leukocytes compared to WT (Fig. 3B), coinciding with the earliest histological evidence of insulitis. At this time point, WT and SKAP2^G153R/G153R^ contained similar proportions of Prdx1^+^ anti-inflammatory (M2-like) macrophages. In contrast, the SKAP2^G153R/G153R^ islets were enriched for transcripts associated with Cxcl9^+^ inflammatory (M1-like) macrophages, dendritic cells, and T and B lymphocytes (Fig. 3B, C). This included increased numbers of islet-infiltrating T regulatory and T follicular helper cells, activated B-cells, and plasma cells, despite the absence of detectable SKAP2 expression in these lymphoid populations. By eight weeks of age, immune infiltration in WT islets approached that observed in SKAP2^G153R/G153R^ animals. However, B lymphocytes remained disproportionately enriched in the *Skap2* mutant animals (Fig. 3C). These transcriptional findings closely mirrored the histologic analyses, which showed accelerated myeloid-dominant insulitis in SKAP2^G153R/G153R^ pancreatic islets, followed by a more robust B cell response at later time points (Fig. S2A).

**Figure 3.**
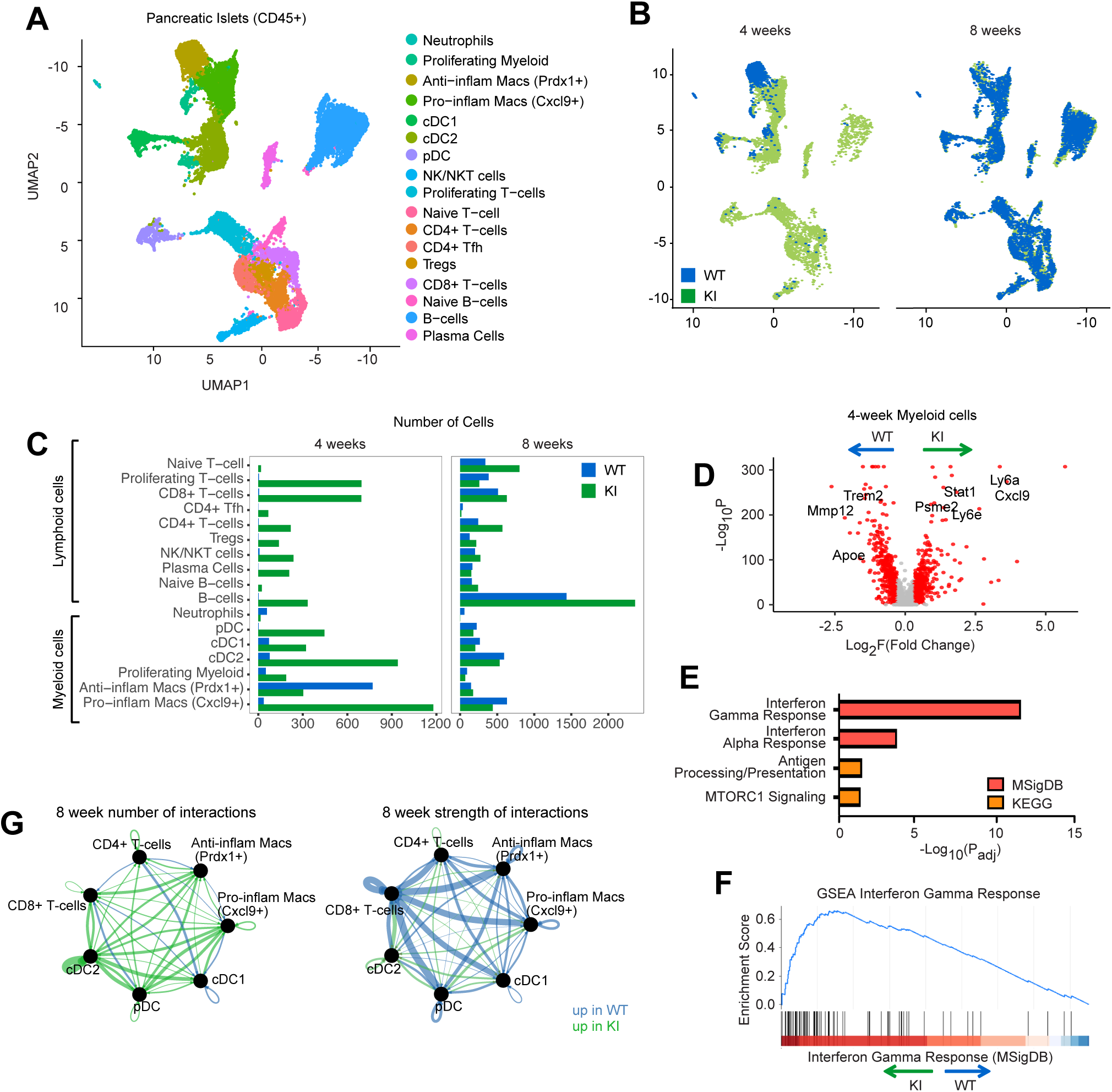
SKAP2^G153R/G153R^ mice have early CD45^+^ infiltration into their islets and higher expression of genes involved in IFNγ-driven inflammatory response. **(A)** CD45^+^ cells were sorted from isolated pancreatic islets of female WT or SKAP2^G153R/G153R^ (labeled as KI) mice and 10x single cell sequenced (four-week-old mice n=3, eight-week-old mice n=2). All cells are displayed in a UMAP with individual clusters identifying the indicated cell types. **(B)** Four-week (left) and eight-week (right) CD45^+^ cells with colors representing WT (blue) and KI (green) to compare the populations present in the genotypes over time. **(C)** Bar graphs of the cell populations present at four weeks and eight weeks in the two genotypes. **(D)** Gene set analysis (GSEA) was performed on the myeloid subset isolated from the islets of four-week-old WT and KI mice, and the results were plotted in a volcano plot. Red dots indicate genes showing log_2_ fold difference > 0.3 and -log_10_ > 1.3 (giving a p<0.05), which are default cutoffs in Seurat/Volcano packages. **(E)** MSigDB Hallmark gene set analysis on the most differentially expressed genes in subset myeloid cells from the islets of WT versus SKAP2^G153R/G153R^ (KI) animals. **(F)** A gene enrichment plot of the IFNγ response transcripts, which was the highest expressed gene set in the SKAP2^G153R/G153R^ (KI) myeloid cells, showing significant enrichment of IFNγ induced transcripts in the SKAP2^G153R/G153R^ cells. **(G)** Graphical representation of cellular interactions between islet immune cells in WT versus SKAP2^G153R/G153R^ (KI), taken from samples of 8-week-old mice, as predicted by CellChat. Data are shown as a ball plot, with line widths indicating predicted numbers and the strength of cellular interactions.

To define the inflammatory programs underlying these differences, we performed gene set enrichment analysis (GSEA) (Maleki et al., 2020) on differentially expressed genes in islet-infiltrating immune cells. At four weeks of age, the myeloid cells in SKAP2^G153R/G153R^ mice expressed significantly higher levels of proinflammatory genes, including *Cxcl9, Ly6a,* and *Ly6e,* compared to WT controls (Fig. 3D). This transcriptional difference diminished by eight weeks of age (Fig. S4A). Hallmark gene set analysis (MSigDB) (Liberzon et al., 2015) of the differentially expressed genes in four-week islet macrophages revealed a dominant IFNγ-stimulated transcriptional signature, with a more modest contribution from type I IFN-responsive genes (Fig. 3E). Enrichment of IFNγ response genes was strongest in SKAP2^G153R/G153R^ myeloid cells (Fig. 3F).

We next examined predicted intercellular communication networks using CellChat v2 (Jin et al., 2025) in eight-week-old immune populations. SKAP2^G153R/G153R^ immune cells exhibited an increased number of predicted ligand-receptor interactions across immune subsets compared with WT controls (Fig. 3G and Fig. S4B). However, the expected interaction strength was reduced overall compared to WT, suggesting that the immune cells in SKAP2^G153R/G153R^ islets are broadly primed for interaction but may engage in more transient or less stable signaling contacts.

To examine transcript levels for specific activation markers, we conducted a subset analysis of plasmacytoid dendritic cells (pDCs), conventional dendritic cells (cDCs), and pro- and anti-inflammatory macrophages. At four weeks of age, cDCs from the islets of SKAP2^G153R/G153R^ mice had higher transcript counts for *Cd80* and *Cd86*, as did the inflammatory macrophages. *Cd80* gene expression remained elevated in islet inflammatory macrophages at eight weeks of age. In addition, cDCs at four weeks had significant upregulation of *Cd274* (PD-L1), consistent with our spectral flow (Fig. S3E). By eight weeks of age, *Cd274* expression was similar between WT and SKAP2^G153R/G153R^ cDCs (Fig. S4C). In contrast, anti-inflammatory macrophages from SKAP2^G153R/G153R^ mice exhibited reduced *Mmp12* expression compared to WT cells. At eight weeks of age, the DC subsets from SKAP2^G153R/G153R^ mice also showed slightly higher levels of MHC class II transcripts.

Together, these data suggest that the enhanced inflammatory insulitis in SKAP2^G153R/G153R^ knock-in mice is initiated by proinflammatory, IFNγ-responding macrophages and dendritic cells, consistent with the fact that SKAP2 is predominantly expressed in myeloid cells.

### SKAP2 G153R mutation enhances dendritic cell-mediated T cell activation

To investigate how hyperactivated SKAP2-mediated integrin signaling in myeloid cells could accelerate autoimmune T1D in NOD mice, we followed the observation that dendritic cells lacking SKAP2 exhibit reduced adhesion to T cells and impaired antigen-specific T cell priming (Reinhold et al., 2009). As the SKAP2 G153R variant is a gain-of-function mutation that increases myeloid cell adhesion (in human cells from the index patient) (Rutsch et al., 2021), we hypothesized that dendritic cells from SKAP2^G153R/G153R^ mice would display enhanced capacity to engage and activate antigen-specific T cells. To test this hypothesis *in vivo*, diabetogenic chromogranin A-specific CD4^+^ T cells from NOD BDC2.5 transgenic mice (BDC2.5 T cells) were isolated by fluorescence-activated cell sorting, labeled with carboxyfluorescein succinimidyl ester (CFSE), and adoptively transferred into WT or SKAP2^G153R/G153R^ recipient mice. T cell proliferation was evaluated 72 hours after cell transfer by flow cytometric analysis of CFSE dilution in cells collected from the spleen, pancreatic lymph nodes, and control lymph nodes (axillary, branchial, and inguinal) (Fig. 4A). BDC2.5 T cells collected from the pancreatic lymph nodes of SKAP2^G153R/G153R^ recipients exhibited significantly increased proliferation compared with those recovered from WT recipients (Fig. 4B, C). In contrast, no differences in proliferation were observed in the spleen or control lymph nodes (Fig. 4C), consistent with the absence of chromogranin A protein expression in these tissues. These findings suggest that antigen-presenting cells in SKAP2^G153R/G153R^ mice prime chromogranin A-specific T cells more efficiently *in vivo*.

**Figure 4.**
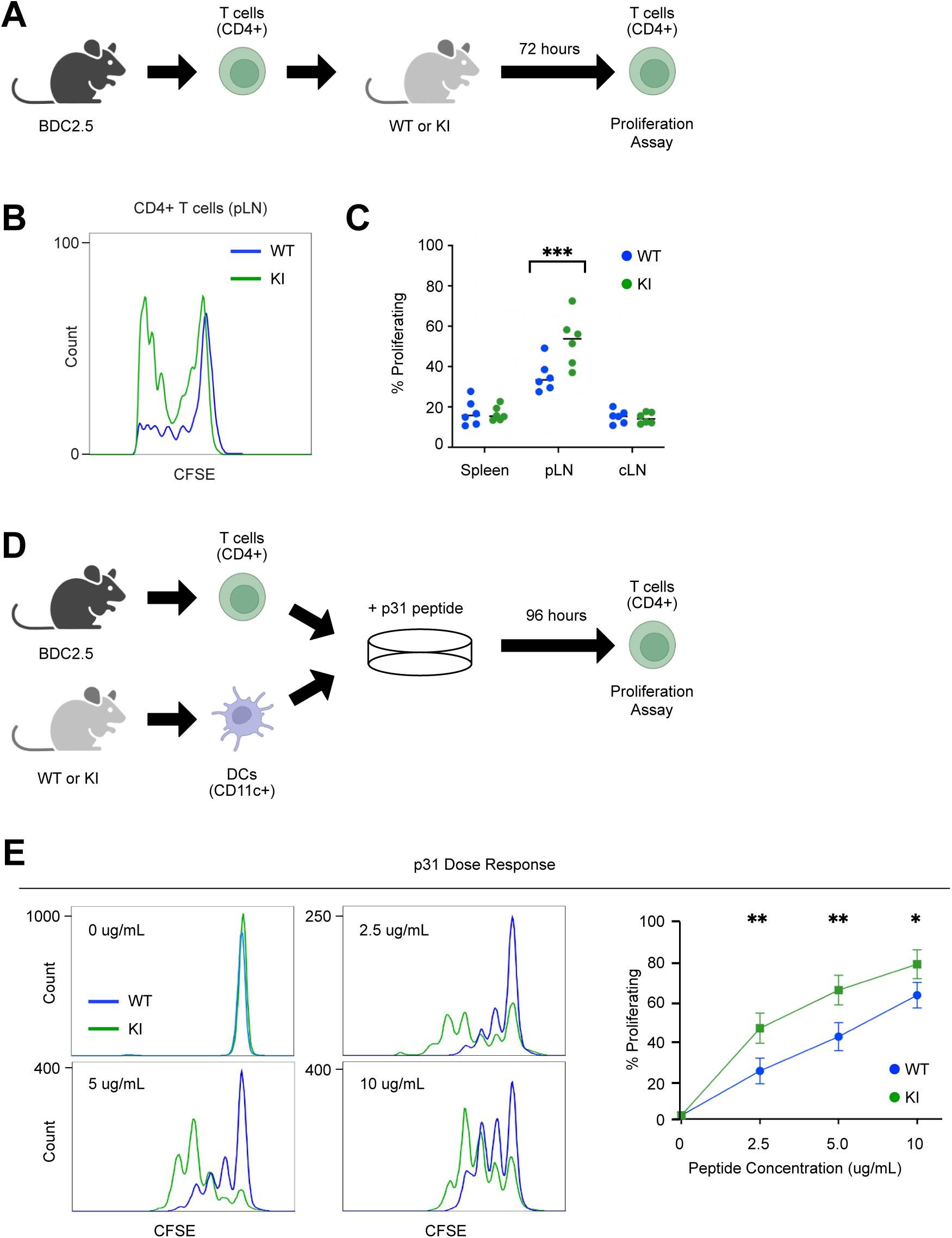
Skap2^G153R/G153R^ dendritic cells have an increased ability to induce T cell proliferation. **(A)** Graphic of the experiment; CD4^+^ T cells isolated from the spleen and control LNs (cervical, inguinal, axillary, and brachial) of BDC2.5 mice were adoptively transferred into 4-week-old female WT or SKAP2^G153R/G153R^ (labeled as KI) recipients. The spleen, pancreatic lymph nodes (pLN), and control lymph nodes (cLN) from the recipient mice were isolated, and the proliferation of the donor CD4^+^ T cells was evaluated by FACs analysis of CFSE dilution in proliferation graphs (n=6 per group). **(B)** Representative proliferation plots comparing CD4^+^ BDC2.5 T cells after being injected into WT or KI mice. **(C)** Quantitation of proliferation data. The data are representative of two independent experiments comparing the spleen, pLN and cLN. **(D)** Graphic of the experiment: dendritic cells from WT or KI mice and CSFE-labeled T cells from BDC2.5 mice were isolated, then co-cultured in the presence of increasing amounts of p31 peptide for 96 hours, after which T cell proliferation was assessed by CFSE dye dilution. **(E)** p31 peptide was added at concentrations of 0, 2.5, 5, and 10 μg/uL in both WT and KI co-culture, then the percent BDC2.5 cell proliferation was determined (n=3 per concentration). Representative flow cytometry plots from the four peptide concentrations and quantification of the experiment are shown. Data are representative of three independent experiments. P values were calculated by two-way ANOVA; *=p<0.05; **=p<0.01; ***=p<0.001.

To determine whether this effect reflected a cell-intrinsic enhancement of T cell priming, we performed *in vitro* co-culture studies. Equal numbers of dendritic cells isolated from WT or SKAP2^G153R/G153R^ mice were cultured with CFSE-labeled BDC2.5 T cells in the presence of increasing concentrations of p31 peptide, the chromogranin A antigenic epitope recognized by BDC2.5 T cells (Nikoopour et al., 2011). After 96 hours T cell proliferation by was quantified by flow cytometry (Fig. 4D). As expected, BDC2.5 T cells did not proliferate in the absence of the p31 peptide, confirming that neither WT nor SKAP2^G153R/G153R^ dendritic cells activated diabetogenic T cells in antigen-independent manner (Fig. 4E). in the presence of p31 peptide, dendritic cells from SKAP2^G153R/G153R^ mice induced significantly higher proliferation of BDC2.5 T cells than WT dendritic cells at all concentrations of peptide tested (Fig. 4E). Together, these data demonstrate that the SKAP2 G153R mutation enhances the intrinsic capacity of dendritic cells to prime and activate antigen-specific T cells responses, providing a mechanistic link between dysregulated myeloid integrin signaling and accelerated autoimmune diabetes.

### SKAP2^G153R/G153R^ dendritic cells display increased adhesion to antigen-specific T cells

Given that SKAP2 functions within the integrin signaling pathway, we postulated that the enhanced ability of SKAP2^G153R/G153R^ dendritic cells to prime antigen-specific T cell responses reflects increased adhesive interactions (i.e., conjugate formation) with the T cells. To test this, dendritic cell/T cell conjugate formation was assessed by co-culturing dendritic cells isolated from WT or SKAP2^G153R/G153R^ mice with BDC2.5 T cells in the presence of increasing concentrations of p31 peptide for 30 minutes at 37°C, followed by quantification of conjugates by flow cytometry (Fig. 5A). SKAP2^G153R/G153R^ dendritic cells formed significantly more conjugates with BDC2.5 T cells than WT dendritic cells, even in the absence of p31 peptide. The presence of p31 peptide further increased conjugate formation by SKAP2^G153R/G153R^ dendritic cells relative to WT controls (Fig. 5B), indicating that the SKAP2 G153R mutation enhances dendritic cell / T cell independently of antigen recognition, which is augmented upon antigen engagement. To characterize the dynamics of dendritic cell/T cell interactions, we used time-lapse confocal microscopy to quantify the number and duration of immunological synapses formed in the presence of the p31 peptide (Fig. 5C-E). Consistent with the conjugate assay, individual SKAP2^G153R/G153R^ dendritic cells engaged a greater number of antigen-specific BDC2.5 T cells over the 2-hour co-culture period. Unexpectedly, the duration of individual dendritic / T cell interactions was significantly shorter for SKAP2^G153R/G153R^ dendritic cells compared to WT cells. This observation closely paralleled predictions from CellChat analysis of the scRNAseq data set (Fig. 3G), which indicated increased numbers of intercellular interactions with reduced interaction strength in SKAP2^G153R/G153R^ immune cell populations. Together, these findings suggest that SKAP2^G153R/G153R^ dendritic cells form more frequent but shorter-lived contacts, consistent with enhanced integrin-dependent adhesion, enabling rapid and efficient antigen presentation to multiple T cells over time.

**Figure 5.**
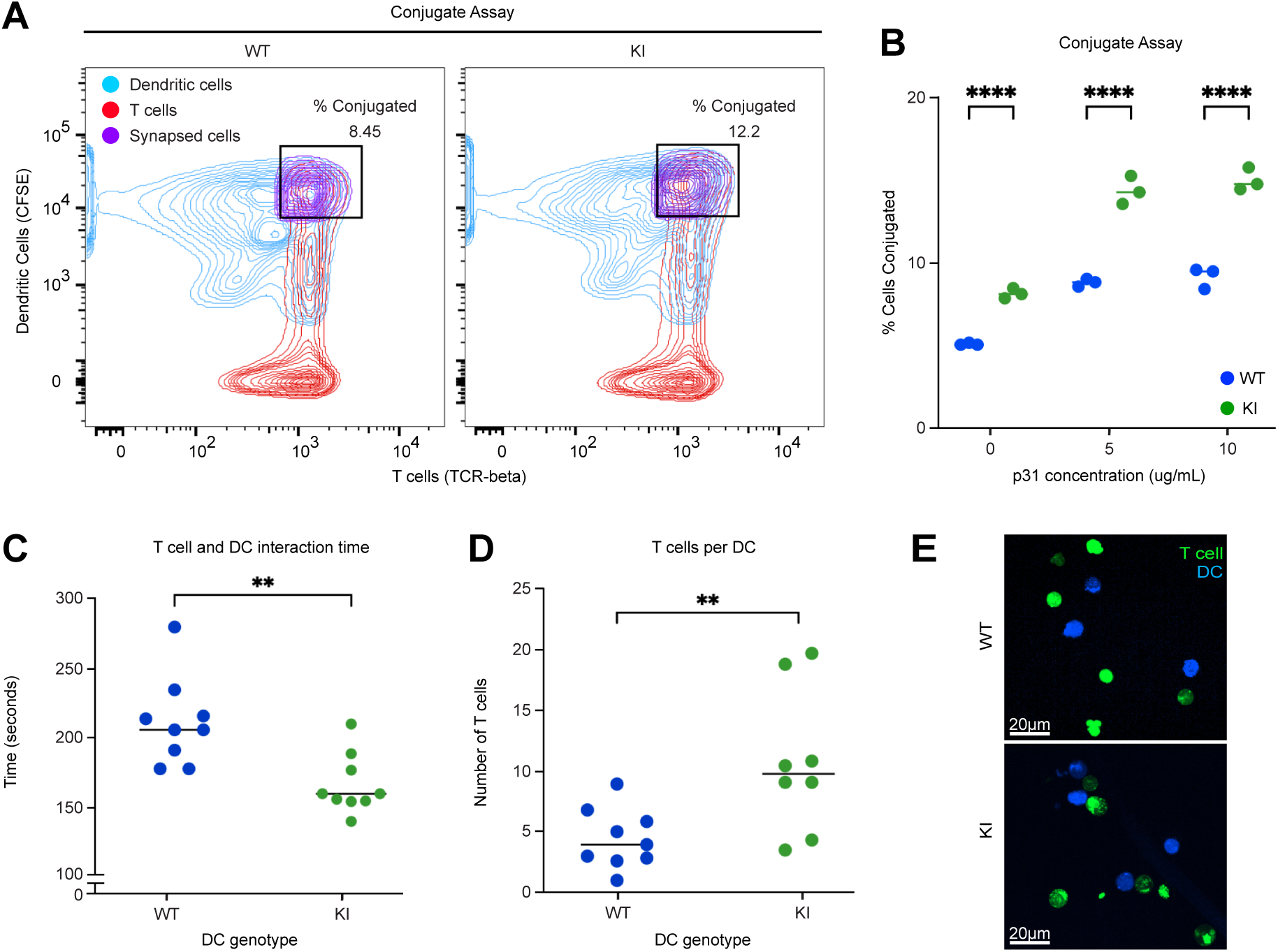
SKAP2^G153R/G153R^ dendritic cells have greater adhesion to antigen-specific T cells. **(A,B)** CSFE-labelled dendritic cells from female WT or SKAP2^G153R/G153R^ (labeled as KI) mice and T cells from BDC2.5 mice were isolated and co-cultured with 0-10 μg/μL of p31 peptide for 30 minutes at 37°C, then analyzed by FACs to detect doublets formed by conjugated dendritic cells and T cells. The percent of conjugated cells was quantified at each peptide concentration (n=3). **(C-E)** BDC2.5 T cells (labeled with cell trace violet) and WT or KI dendritic cells (labeled with CSFE and pre-loaded with 5 μg/μL of p31 peptide) were cultured together and visualized using confocal microscopy over a 2-hour time period. Interaction duration **(C)** and the number of T cells associated with a given dendritic cell **(D)** over the entire co-culture time were quantified using Imaris software. **(E)** Representative photos of the experiment. Data are representative of three independent experiments. P values were calculated by two-way ANOVA; **=p<0.01; ****=p<0.0001.

### Hyper-activation of integrin pathways in SKAP2^G153R/G153R^ macrophages and neutrophils

Previous studies of SKAP2 function in integrin signaling have primarily focused on macrophages and neutrophils from *Skap2^-/-^* mice on the C57Bl/6 genetic background. In these experiments, in which SKAP2 deficiency results in impaired macrophage migration, defective formation of membrane protrusions, and reduced integrin-dependent neutrophil oxidative burst (Alenghat et al., 2012; Boras et al., 2017; Swanson et al., 2008). Based on these loss-of-function phenotypes, we postulated that the SKAP2 G153R gain-of-function mutation would produce opposite effects in these myeloid cell populations. To test this, we assessed macrophage migration using a scratch wound assay, in which confluent monolayers of bone marrow-derived macrophages from WT or SKAP2^G153R/G153R^ mice were mechanically wounded and cell migration into the wounded area was examined over time. SKAP2^G153R/G153R^ macrophages migrated much more rapidly than the WT control cells (Fig. 6A, B). To directly explore integrin-dependent actin remodeling, we used a bead-based anti-α_v_ integrin antibody crosslinking assay previously used to study *Skap2^-/-^*macrophages (Alenghat et al., 2012). Macrophages were stimulated with uncoated or anti-α_v_ antibody-coated polystyrene beads, followed by fixation and staining with anti-SKAP2 and phalloidin to visualize polymerized F-actin for confocal imaging (Fig. 6C, D). In response to integrin crosslinking, SKAP2^G153R/G153R^ macrophages exhibited increased membrane ruffling and a higher frequency of filopodial extensions compared to WT control cells, consistent with a stronger integrin signaling response.

**Figure 6.**
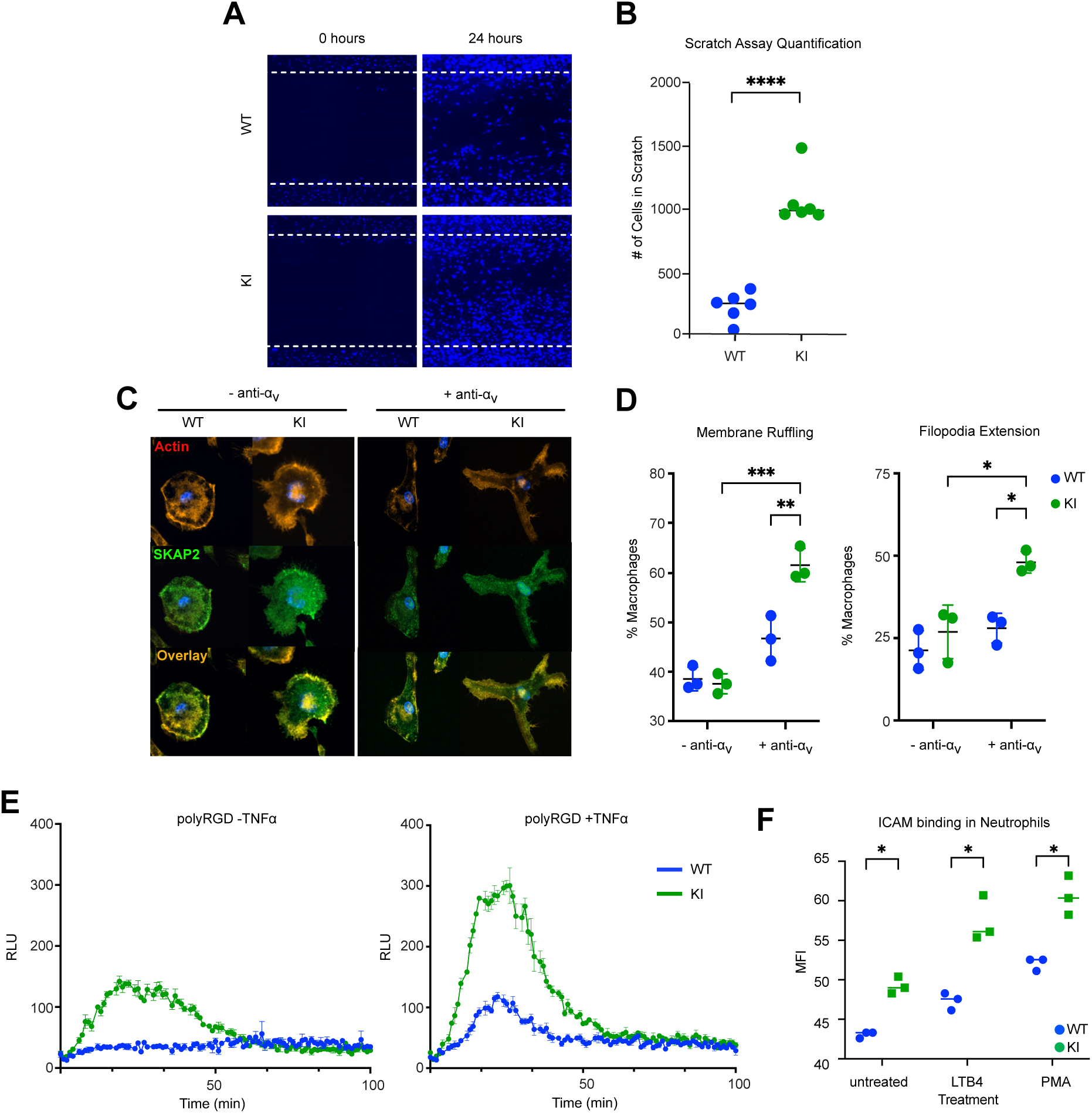
Bone marrow-derived myeloid cells from SKAP2^G153R/G153R^ mice have hyperactive integrin signaling. **(A-B)** Bone marrow-derived macrophages (BMDM) from female WT or SKAP2^G153R/G153R^ (KI) mice were expanded on tissue culture plates to form confluent monolayers, then scratched with a P200 pipette tip. Macrophage migration into the scratch was monitored for 24 hours. Cells were fixed and stained with DAPI (n=6). The images were quantified by cell counting using QuPath’s “cell detection” function and confirmed by manual counting. **(C)** Confocal images of WT or KI BMDMs plated on glass coverslips, then stimulated with uncoated polystyrene beads (as controls) or with beads coated in anti-α_v_ integrin antibody as an integrin stimulus. After 20 mins, cells were fixed and then stained for F-actin with phalloidin (red) and anti-SKAP2 (green). Merged images show the overlap of the stains in yellow. **(D)** Quantification of BMDMs with membrane ruffling (colocalization of F-actin and SKAP2 at the cell periphery) and filopodia extensions (5 or more filopodia) (n=3). **(E)** Bone marrow-derived neutrophils from WT or KI mice were plated on poly-RGD-coated wells with no stimulation or TNFα in the presence of isoluminol, then placed on a luminescence reader for 100 minutes to monitor neutrophil ROS production. Data are representative of three independent experiments; cells were pooled from three mice per experiment. **(F)** Bone marrow cells from WT and KI mice were incubated for 10 min at room temperature with recombinant ICAM-1/Fc, then stimulated with LTB4 or PMA, then counterstained for the human Fc domain on the ICAM/Fc reagent and analyzed by flow cytometry. Dots show values from separate animals. P values were calculated by unpaired t-tests. Data represent the difference between means ± SEM; *=p<0.05; **=p<0.01; ***=p<0.001; ****=p<0.0001.

To place these findings in the context of established SKAP2 loss-of-function phenotypes, we performed the same assays in parallel using macrophages from C57BL/6J *Skap2^-/-^* knockout mice. As previously reported (Alenghat et al., 2012), SKAP2-deficient macrophages showed impaired migration and minimal filopodial formation following integrin engagement, while SKAP2^G153R/G153R^ macrophages displayed enhanced migratory and cytoskeletal responses relative to WT controls (Fig. S5A, B). Notably, the heightened migratory phenotype of SKAP2^G153R/G153R^ macrophages closely mirrors that of monocyte-derived macrophages from the index T1D patient harboring the SKAP2 mutation (Rutsch et al., 2021).

Deficiency of SKAP2 also impairs integrin-stimulated reactive oxygen species (ROS) production in neutrophils (Boras et al., 2017). To examine whether the SKAP2 G153R gain-of-function mutation similarly enhanced neutrophil integrin signaling, bone marrow-derived neutrophils were plated on poly-RGD protein surfaces in the absence or presence of TNFα to enhance integrin responses (Abram et al., 2013), and neutrophil ROS production was quantified. In both conditions, SKAP2^G153R/G153R^ neutrophils displayed significantly increased ROS release compared to WT control neutrophils following integrin engagement (Fig. 6E), consistent with hyperactive integrin signaling. Integrin signaling in neutrophils requires activation of Bruton’s Tyrosine Kinase (BTK) upstream of SKAP2, which can be pharmacologically inhibited by ibrutinib (Volmering et al., 2016). Treatment with ibrutinib attenuated the ROS response of SKAP2^G153R/G153R^ neutrophils in a dose-dependent manner, similar to that observed in WT control cells (Fig. S5C), indicating that the hyperactive signaling caused by the SKAP2 G153R gain-of-function mutation remains dependent on canonical BTK-mediated pathways.

To directly measure integrin activation status on WT versus SKAP2^G153R/G153R^ neutrophils, we used a soluble ICAM1/Fc binding assay to measure the affinity of CD18 integrins (LFA-1 and Mac-1) on the neutrophil surface (Amini et al., 2025). Both resting and stimulated (LTB4 or PMA) SKAP2^G153R/G153R^ neutrophils displayed significantly greater binding of soluble ICAM1/Fc than WT cells (Fig. 6F), demonstrating enhanced integrin activation. Together, these findings establish that the SKAP2 G153R gain-of-function mutation drives heightened integrin activation in neutrophils and downstream functional responses in myeloid cells. This provides mechanistic insight into the amplified inflammatory phenotype of SKAP2^G153R/G153R^ mice.

### SKAP2^G153R/G153R^ mice on the C57BL/6 genetic background develop autoimmunity

All the experiments described above were conducted using mice on the NOD genetic background. To determine whether the SKAP2 G153R gain-of-function mutation predisposes to autoimmunity on a genetically resistant background, we backcrossed the NOD SKAP2^G153R/G153R^ mice into the C57BL/6 background, which is known to be much more resistant to spontaneous autoimmunity (Morel, 2004). Mice harboring the SKAP2^G153R^ knock-in allele were backcrossed for 12 generations, and breeder animals were genotyped by allele-specific PCR to assess genomic background composition. The six breeders examined had 95% C57BL/6 genome content, with the remaining NOD-derived sequences mapping to *Skap2* itself and a region on chromosome 13, not previously associated with diabetes susceptibility (Fig. S6A) (Chen et al., 2018; Thayer et al., 2010). These mice were used for further study.

SKAP2^G253R/G153R^ mice on the C57BL/6 genetic background did not develop spontaneous diabetes and did not manifest overt signs of tissue inflammation as determined by histology. A subset of animals developed modest splenomegaly, although this finding was variable (not shown). To investigate whether the SKAP2 G153R gain-of-function mutation promoted systemic autoimmunity, we analyzed serum samples from cohorts of WT and SKAP2^G153R/G153R^ C57BL/6 mice using PhIP-seq. Surprisingly, both male and female SKAP2^G153R/G153R^ C57BL/6 mice exhibited broad autoantibody reactivity with diverse specificities, beginning as early as 2 months of age. In contrast, autoantibodies in C57BL/6 WT mice were minimal and barely detectable (Fig. 7A). These findings suggest that SKAP2^G53R/G153R^ mice develop low-grade spontaneous systemic autoimmunity on the C57BL/6 genetic background.

**Figure 7.**
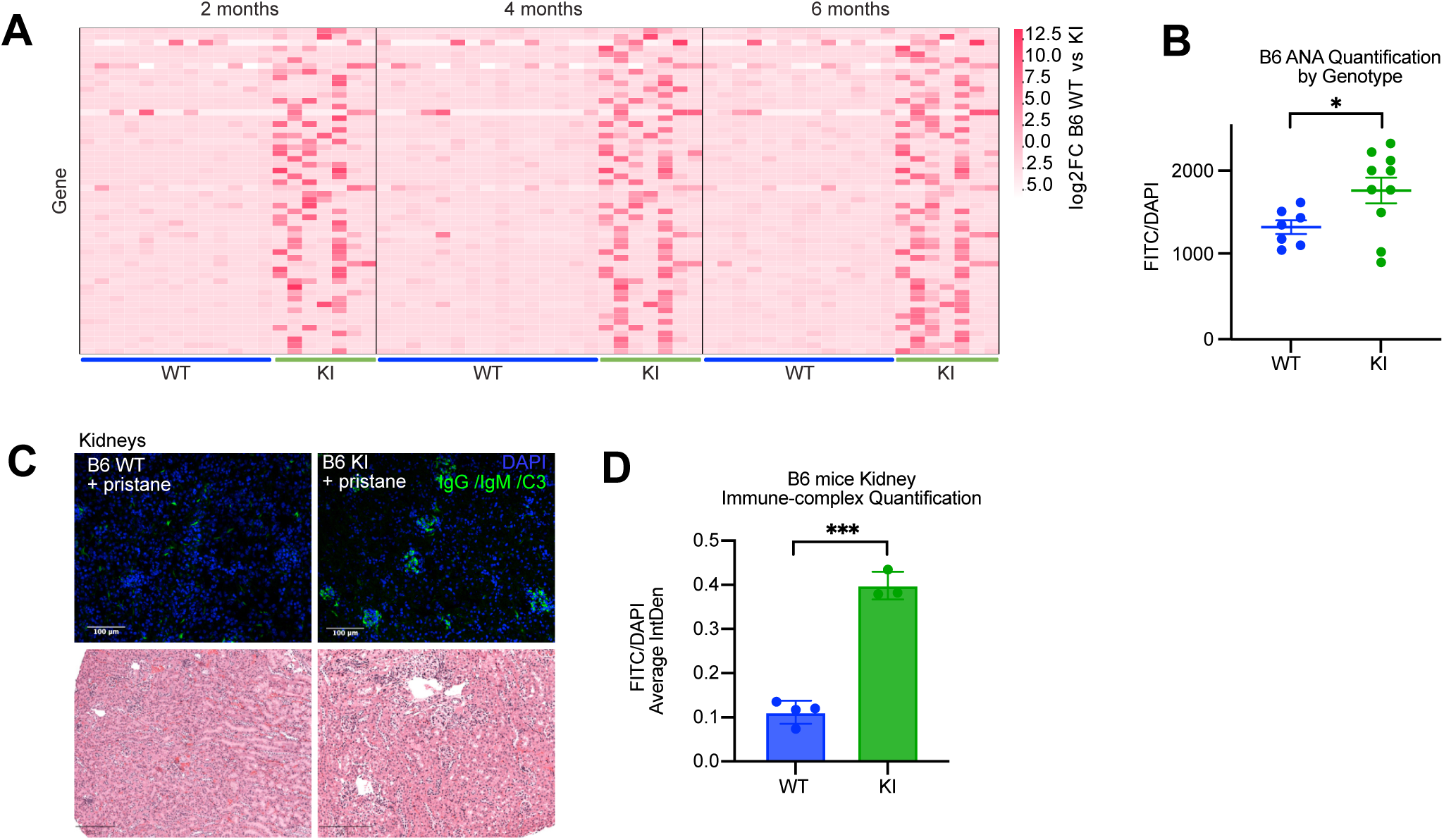
C57BL/6 SKAP2^G153R/G153R^ mice have baseline autoimmunity and show accelerated lupus-like autoimmune disease in the pristane model. **(A**) Serum samples from female C57BL/6 WT (n=13) or B6 SKAP2^G153R/G153R^ (KI) (n=7) mice at ages two, four and six months were analyzed for autoantibodies by PhIP-Seq. Heatmaps showing gene level log₂ fold-change (log₂FC) values across three longitudinal timepoints in the PhIP-Seq study for C57BL/6 WT and B6 SKAP2^G53R/G153R^ mice. Values represent individual sample log₂FCs relative to WT controls. Only genes exhibiting a log₂FC>2 at any timepoint are shown (see Table S3 for a list of antigens displayed). For genes represented by multiple peptides, log₂FC values were averaged to derive a single gene-level enrichment score **(B)** Serum samples from female C57BL/6 WT (n=7) or B6 SKAP2^G153R/G153R^ (KI) (n=10) were taken 2 months after pristane injection and analyzed for anti-nuclear antibody (FITC) plus DAPI nuclear staining on Hep-2 cells. Staining was imaged by fluorescence microscopy, and the ratio of FITC to DAPI nuclear staining was quantified by Image J analysis. **(C, D)** Immune complex deposition in the kidneys of C57BL6 WT and B6 SKAP2^G153R/G153R^ (KI) mice at 2 months following pristane injection was assessed by IgG/IgM/C3 immunofluorescence staining and quantification. Lower panels show standard H+E staining of kidney samples to reveal increased inflammatory cell infiltration in the KI samples. In bar graphs, each dot represents samples from a single mouse. P values were calculated by unpaired t-tests. Data represent the difference between means ± SEM; *=p<0.05; ***=p<0.001.

To evaluate whether the SKAP2 G153R gain-of-function mutation exacerbates autoimmune disease under inflammatory challenge, we employed the pristane-induced model of lupus-like autoimmunity (Freitas et al., 2017; Zhou et al., 2025). Serum samples from mixed-sex cohorts of WT and SKAP2^G153R/G153R^ C57BL/6 mice were analyzed at two months following pristane injection for anti-nuclear antibody (ANA) by immunofluorescence. SKAP2^G153R/G153R^ C57BL/6 mice exhibited significantly higher levels of ANA compared to WT controls (Fig. 7B). This enhanced autoantibody response was associated with early-onset immune complex-mediated glomerulonephritis in SKAP2^G153R/G153R^ C57BL/6 mice, as evidenced by histologic examination of kidney sections showing immunoglobulin and complement deposition (Fig. 7C, D). Three months following pristane injection, SKAP2^G153R/G153R^ C57BL/6 mice tended to have larger spleens with higher myeloid cell counts but lower T cell numbers than WT controls (Fig. S6B). The SKAP2^G153R/G153R^ C57BL/6 mice also displayed obvious accelerated formation of peritoneal lipogranulomas, which formed masses on the stomach and peritoneum (Fig. S6C). Together, these findings demonstrate that the SKAP2 G153R gain-of-function mutation promotes the development of autoimmunity even in genetically resistant C57BL6 mice.

## DISCUSSION

The genetic susceptibility to type 1 diabetes (T1D) is complex, with variants from many immune regulatory genes contributing to disease risk. Variants in the gene encoding the integrin signaling adapter protein, *SKAP2*, are among the most common single-nucleotide polymorphisms associated with T1D (Robertson et al., 2021). We identified a single individual with a complex form of T1D (diabetes associated with other autoimmune manifestations) who carried a *de novo* mutation in one *SKAP2* allele (Rutsch et al., 2021). Functional studies of this single amino acid substitution (G153R) suggested that it produced a constitutively activated form of SKAP2, leading to hyperactivation of integrin signaling pathways and resulting in more adhesive and migratory myeloid leukocytes (Rutsch et al., 2021). To define the mechanisms by which dysregulated integrin signaling in myeloid cells could lead to T1D, we generated mice with the same single amino acid substitution, SKAP2 G153R, as was mutated in the patient. By making this point mutation in the diabetes-susceptible NOD strain of mice, we were able to study the effect on T1D progression. SKAP2 G153R mutant mice, particularly homozygous SKAP2^G153R/G153R^ female mice, exhibited accelerated inflammatory insulitis, resulting in more rapid development of frank T1D in ∼90% of animals. The disease was characterized by early invasion of myeloid cells (macrophages and dendritic cells) into pancreatic islets, with these cells exhibiting an IFN-γ-driven inflammatory transcriptional signature. These myeloid cells are likely responding to cytokines produced by islet antigen-specific T cells, which have been activated by SKAP2^G153R/G153R^ dendritic cells. This inflammatory insulitis likely leads to the rapid breakdown of β cells, causing the release of islet peptides that accelerate the formation of islet (and broadly reactive) T cells, due to the increased ability of SKAP2^G153R/G153R^ dendritic cells to prime T-cell responses. As a result, SKAP2^G153R/G153R^ mice develop a broad spectrum of self-reactive autoantibodies and progressive immune-mediated destruction of islet cells, leading to the onset of frank T1D. The increased antigen-presenting capacity of SKAP2^G153R/G153R^ dendritic cells is likely due to increased dendritic cell/T cell conjugate formation. Hyperactive integrin signaling in macrophages and neutrophils likely contributes to the disease’s inflammatory nature. Indeed, SKAP2^G153R/G153R^ neutrophils display higher levels of activated integrins even in resting cells. These phenotypes closely mirror the disease state in the index SKAP2 G153R patient, demonstrating the model’s physiologic relevance.

The significance of this study lies in the fact that single-nucleotide polymorphisms in *SKAP2* are frequently observed in T1D individuals, even though the mutation we are studying, SKAP2 G153R, is rare and has not been previously reported in patients with autoimmune disease (Rutsch et al., 2021). How the other *SKAP2* polymorphisms contribute to risk for T1D or other autoimmune/inflammatory diseases remains unclear. Indeed, many of these polymorphisms are in non-coding regions of *SKAP2*. One of the better-studied *SKAP2* polymorphisms associated with T1D, rs7804356, is in the third intron of *SKAP2* and is associated with reduced *SKAP2* RNA expression in phorbol ester-stimulated Epstein-Barr Virus (EBV)-transformed B lymphocytes (Floyel et al., 2021; Ram et al., 2016). Interestingly, this polymorphism has also been associated with rheumatoid arthritis in a Pakistani patient cohort (Kiani et al., 2015). A recent study suggested that reduced SKAP2 expression may increase pancreatic cell susceptibility to the apoptotic effects of proinflammatory cytokines. However, the molecular mechanisms underlying this phenomenon remain unclear (Floyel et al., 2021). Given that *Skap2^-/-^* mice have no reported alterations in glucose homeostasis (data not shown) and instead exhibit a phenotype consistent with immunodeficiency rather than autoimmunity, it is hard to speculate on how reduced SKAP2 expression may confer genetic risk for T1D. Future studies are needed to determine how non-coding *SKAP2* polymorphisms impact integrin signaling in myeloid cells.

We focused on dendritic cell priming of T cell reactivity in SKAP2^G153R/G153R^ mice as a primary driver of autoimmune insulitis. However, it seems likely that hyperactive integrin signaling in macrophages and neutrophils may also contribute to the inflammatory phenotype in SKAP2^G153R/G153R^ mice. SKAP2 has been implicated in the regulation of TLR4 signaling, in part by associating with the tyrosine phosphatase SHP-1 (Takagane et al., 2022). Thus, an increased migratory phenotype in macrophages may contribute to heightened inflammatory responses, while the heightened ROS release by SKAP2^G153R/G153R^ neutrophils would contribute to tissue damage. Ultimately, mice with lineage-specific expression of SKAP2 G153R will need to be generated to determine the relative importance of dendritic cells, macrophages, and neutrophils to the T1D phenotype.

We also found that the SKAP2 G153R mutation led to enhanced autoimmunity in C57BL/6 mice. This suggests that the increased integrin signaling caused by the SKAP2 mutation may affect other disease models, such as colitis or antibody-induced arthritis. SKAP2 is also highly expressed in eosinophils (Immgen), which may lead to increased allergic inflammation in different disease settings – allergy is also a characteristic of the index SKAP2 G153R patient (Rutsch et al., 2021). These remain future avenues of research using this animal model.

An unanticipated finding in this study was that, while p31 peptide-loaded SKAP2^G153R/G153R^ dendritic cells bound more antigen-specific BDC2.5 T cells than WT dendritic cells, the SKAP2^G153R/G153R^ dendritic cells bound the T cells for a shorter time. Interestingly, the CellChat analysis of our scRNAseq data predicted just this – more intracellular interactions in SKAP2 knock-in immune cells, but with weaker interaction strengths than in WT immune cells. It is well recognized that establishing the immunologic synapse during antigen presentation between dendritic cells and T cells involves initiating intracellular signaling in both cell types – it is a bilateral response (reviewed in Martin-Cofreces et al., 2018). The duration of dendritic T-cell synapses is affected by peptide affinity and dose, both of which contribute to TCR signaling strength. The duration of stable dendritic cell/T cell interactions is estimated to be 3–5 hours, after which a detachment phase occurs, likely due to downregulation of T cell adhesion receptors (Beltman et al., 2009). The effect of increased adhesive function on the dendritic cell side of the immunologic synapse remains to be explored. It is possible that the increased actin dynamics observed in SKAP2^G153R/G153R^ dendritic cells may lead to more rapid surface recycling of integrin receptors, resulting in more frequent but shorter synapse formation with antigen-specific T cells. Attempts to manipulate antigen-presenting cell signaling responses presents a new avenue for modulating T cell-mediated immune responses.

Several mutations in integrin signaling molecules have been described, leading to the development of autoinflammation and immune dysfunction (Papa et al., 2020). Many of these mutations affect actin cytoskeletal dynamics and, as a group, are referred to as “actinopathies”. While most are loss-of-function mutations, some are also gain-of-function mutations, including variants in WASP, which directly associate with SKAP2. Given the dramatic effect of the SKAP2 G153R mutation on actin polymerization in macrophages from the knock-in mice and in monocytes from the index patient, it is likely that the SKAP2 G153R mutation could also be considered as an “actinopathy” variant.

The fact that ibrutinib completely reversed the hyperactive integrin signaling response of SKAP2^G153R/G153R^ neutrophils suggests a possible therapeutic approach for *SKAP2*-associated T1D cases. Indeed, several tyrosine kinase inhibitors have been shown to block diabetes in the NOD model (though ibrutinib has not been tested) (Louvet et al., 2008; Zeng et al., 2022). The modulation of leukocyte integrin function by blocking monoclonal antibodies or inhibiting intracellular signaling is being developed and used in various immune-mediated disease settings. Similarly, our exome sequencing efforts in T1D patients have identified additional individuals and families with polymorphisms in other integrin signaling molecules, including a multigenerational family with dominant inheritance of a *SKAP2* mutation in the N-terminal dimerization domain, which is predicted to function similarly to the G153R substitution. This suggests that alterations in integrin signaling may contribute broadly to the genetic risk of T1D. Further study of these pathways in diabetes is needed to justify testing of integrin-blocking approaches in individuals at high risk for type 1 diabetes.

## MATERIALS AND METHODS

### Mice

The single-nucleotide variant c.605G>A; p.G153R of murine *Skap2* (NM_018773) was introduced using CRISPR/Cas9 genome editing by the Genetically Engineered Murine Models (GEMM) Core at the University of Colorado (Robinson et al., 2023). CRISPOR and the Broad Institute sgRNA Design software were used to design guide RNAs (Doench et al., 2016; Haeussler et al., 2016). The guide RNA with target sequence TATTATTACGGGAGCGATAAAGG, which was predicted to cut 5 base pairs downstream of G153, was selected, and then its activity was verified by incubating it along with Cas9 protein (Integrated DNA Technologies, Alt-R *Streptococcus pyogenes* HiFi Cas9 Nuclease V3) and a PCR product containing the target sequence, then comparing the ratio of cut to uncut PCR product. *In vitro* fertilization was performed to generate NOD/ShiLtJ (NOD) zygotes, which were injected with the above guide RNA (5ng/ml), Cas9 protein (20ng/ml), and DNA repair template (25 5ng/ml). The HDR repair template spanned 162 nucleotides, with the G>A point mutation in the center. Three additional silent mutations were introduced into the HDR template to avoid the Cas9 cutting the desired, modified allele. Zygotes were then transferred into pseudopregnant recipients. F_0_ pups were genotyped by PCR using primers outside the region modified to identify putative positive founders with the correct G>A substitution - out of 27 pups born, 14 showed evidence of homologous repair. Founders were then crossed to WT NOD mice, and F_1_ pups heterozygous for the *Skap2* point mutation were identified and then transferred to the University of California, San Francisco (UCSF). Initial blood glucose and immune cell evaluations were performed on two separate founder lines, which were found to be equivalent. A single founder was selected and backcrossed to WT NOD mice for 5 generations. All subsequent experiments were done with this line. To generate SKAP2^G153R/G153R^ C57BL/6, we backcrossed the NOD variant with C57BL/6 mice for 12 generations and submitted tails for MiniMUGA Background Analysis (Transnetyx) to characterize the strain. BDC2.5 NOD transgenic mice and *Rag2^-/-^* NOD mice are as described (Warshauer et al., 2021). Mice were maintained in the UCSF specific pathogen-free animal facility in accordance with the guidelines established by the Institutional Animal Care and Use Committee and Laboratory Animal Resource Center, and all experimental procedures were approved by the Laboratory Animal Resource Center at UCSF.

### Western blot analysis of sorted immune populations and immunoprecipitation

Monocytes, dendritic cells, neutrophils, B cells, and T cells were isolated from the bone marrow and spleen of WT and SKAP2^G153R/G153R^ knock-in mouse models using FACs sorting to identify cells using antibodies to CD11b, CD11c, Ly6G, CD19, and CD3, respectively (see Table S4 for antibodies used). Cells were directly lysed in Laemmli’s sample buffer, heated to 95°C for 10 min, and then stored at -20°C. Samples were analyzed using 8% SDS-PAGE, transferred to PVDF membrane, blocked with TBST + 1% BSA, then probed with anti-SKAP2 (PA5-92972; Thermofisher) and ERK2 (sc1647; Santa Cruz) antibodies and incubated at 4°C overnight. Blots were washed with TBST and stained with the following secondary antibodies: IRDye® 680RD goat anti-rabbit IgG (926-68071, LI-COR) and IRDye® 800CW goat anti-mouse IgG (926-32210, LI-COR). Blots were washed and scanned using the LI-COR Odyssey CLx. Total bone marrow cells were harvested from WT and SKAP2^G153R/G153R^ mice, then whole cell lysates were prepared as above, or cells were lysed in RIPA buffer containing protease (058929001, Roche) and phosphatase inhibitor (04906845001, Roche) cocktail tablets. Lysates were analyzed as above, using Abs above, as well as an anti-phosphotyrosine antibody (anti-PTyr, 4G10, Invitrogen). For the SKAP2 immunoprecipitation, bone marrow cell lysates were precleared with protein A/G agarose beads (sc-2003, Santa Cruz Biotechnology) for 1 hour at 4°C, with rotation. Pre-cleared lysates were incubated with anti-SKAP2 antibody overnight at 4°C. Protein A/G agarose beads were then added to each sample and incubated for 3 hours at 4°C to capture the antibody-protein complexes. Beads were washed three times with lysis buffer, and bound proteins were eluted by boiling in SDS sample buffer. Eluted proteins were separated by SDS-PAGE, transferred to nitrocellulose membranes, and probed with anti-phosphotyrosine and anti-SKAP2 antibodies as above.

### Blood glucose determination

Nonfasting blood glucose levels in recipient mice were monitored weekly by using an Accu-Chek Aviva Plus glucometer (Roche Diagnostic Corp). Measurements were obtained at approximately the same time of day. Diabetes onset was considered to have occurred when nonfasting blood glucose concentration exceeded 200 mg/dl for 2 measurements separated by one week.

### Tissue cell preparation and flow spectrometry

Single-cell suspensions of splenocytes and lymph node cells were prepared by passing the tissues through a 70 μm cell strainer. Bone marrow cell suspensions were isolated by flushing femurs and tibiae with Hanks-balanced salt solution without calcium or magnesium (HBSS) and 10 mM HEPES. Red blood cells were lysed by resuspending the samples in RBC lysis buffer for 5 min at RT, then re-centrifuging and resuspending in HBSS/HEPES. Islets were purified following standard collagenase protocols as described previously (Tang et al., 2004) and dissociated by incubation in enzyme-free cell dissociation buffer (Gibco). Cell count was determined using a Nucleocounter™ (Chemometec). Cells were stained at a concentration of 1.5 x 10^6^/ml. Non-specific binding was blocked using 0.5 μg anti-CD16/32 antibody, and cells were stained with the panel shown in Table S4. Analysis was performed using an Aurora (Cytek) and analyzed using FlowJo (Treestar). Statistical analysis was performed using GraphPad Prism.

### Serum autoantibody arrays

Serum (30 μl) from twelve-week-old female WT and SKAP2^G153R/G153R^ mice was sent to the UT Southwestern MicroArray core(Kramer et al., 2016) for analysis of autoantibodies on the super array 124 antigen panel. Samples were treated with DNAse I, diluted 1:50, and incubated with an autoantigen array printed on a 16-pad FAST slide. The autoantibodies binding to the antigens on the array were detected with Cy3 labeled anti-IgG and then scanned with GenePix® 4400A Microarray Scanner. The images were analyzed using GenePix 7.0 software to generate GPR files. The averaged net fluorescent intensity (NFI) of each autoantigen was normalized to internal controls (IgG or IgM) or no protein to determine signal-to-noise ratio (SNR) for all antigens. Data represent antibody binding score as defined by log2 signal/SNR+1. Data show the IgG autoantibody scores plotted as heat maps.

### PhIP-seq autoantibody analysis

Bioinformatic analysis of the PhIP-seq experiment to identify autoantibody ‘hits’ was carried out using the PhagePy Package for python (https://github.com/h-s-miller/phagepy). Raw next- generation sequencing (NGS) reads were demultiplexed and aligned to a murine proteome peptide library using the Bowtie algorithm (Langmead and Salzberg, 2012). The murine proteome, derived from Rackaityte et al., 2023, was systematically tiled using 62–amino acid peptides (62-mers) with 19–residue overlaps, ensuring contiguous coverage of the entire proteome and yielding a library of 482,672 unique sequences. Aligned reads were normalized to the number of reads per 100,000 (RPK) to account for sequencing depth. To control for non-specific signal, data were further normalized to the mean of negative control samples, including MockIP (beads only) and *Rag2*^−/−^ (no antibody) immunoprecipitations. For denoising, peptides with log₂FC<1 relative to negative controls were excluded. Peptides exhibiting a log₂ fold-change (log₂FC) > 1 relative to WT controls in 25% of the SKAP2^G153R/G153R^ samples were considered putative hits. See Table S2 and S3 for a list of antigen hits.

### Histology and immunohistochemistry

Mouse pancreases were harvested and fixed overnight in 4% fresh paraformaldehyde (PFA) in PBS. Tissues were processed in a Leica Tissue Processor and embedded in paraffin wax. Tissue sections were cut at 5 μm and placed on Superfrost glass slides. For hematoxylin and eosin (H&E) and immunohistochemical (IHC) staining, slides were baked at 70°C to adhere to the glass slides. Tissues were rehydrated through a series of xylene-ethanol solutions. H&E slides were stained with hematoxylin (Fisher finest) for 30 seconds, washed with running water, dipped in 95% EtOH, and then stained with eosin (Millipore Sigma) for 10 seconds. The slides then went through the ethanol-xylene dehydration series, and the coverslips were removed for imaging. After rehydration, IHC slides were placed in preheated antigen retrieval buffer (20 mM Boric Acid, 10 mM Tris, and 1 mM EDTA pH 8.0) and incubated at 95°C for 40 min using a PT module (Thermo Scientific PT Module). The antigen retrieval solution was cooled to 65°C and placed at RT for 30 min. Slides were washed twice for 5 min each with TBS-T, then blocked with 5% Donkey Serum (DS, Sigma) and stained with antibodies overnight, diluted in DS. Antibody information is listed in Table S4. The following day, slides were washed twice with TBS-T for 5 min each, then placed in 3% H_2_O_2_/TBS-T for 15 min. Sections were washed in TBS-T twice and then incubated with anti-rabbit HRP secondary (Signal Stain(R) Boost Cat # 8114S Cell Signaling Tech) for 1 hour. Slides were washed in TBS-T 3 times for 5 min and detected with DAB (Signal Stain(R) DAB Cat #11725S Cell Signaling Tech). After 5 min in DAB the slides were washed with Milli_Q water (Millipore) and stained with Hematoxylin for 30 seconds, followed by 30 seconds in NH_4_OH (1:100) washed with tap water and dehydrated in ethanol-xylene series and coverslipped. H&E and IHC slides were scanned on a Leica Aperio AT2. QuPath software was used to examine tissue sections.

### MIBI staining

Isolated mouse pancreases were fixed for 24 hours in 4% paraformaldehyde in PBS, washed 3 times in PBS, and stored in 70% Ethanol at -20°C until paraffin processing. Tissue was infiltrated with paraffin wax (Leica /ASP300S) and then embedded into paraffin blocks. Paraffin embedded tissue blocks (FFPE) were cut at a thickness of 5mm and mounted onto gold-sputtered microscope slides for Multiplex Ion Beam Imaging processing (IonPath). Tissue Gold Slides were baked at 70°C overnight, and dewaxing and staining were done according to Ionpath protocol. Briefly, baked tissue was deparaffinized, dehydrated, and then antigen retrieved using high pH (Dako Target Retrieval) for 40 min at 97°C, followed by cooling to 65°C in a Lab Vision PT module. (Thermo Fisher Scientific). Slides were cooled to room temperature for 30 min and washed twice in TBS-T (Ionpath). Tissues were blocked with 5% donkey serum (DS, Sigma-Aldrich)-TBS-T for 1 hour at room temperature. The antibody cocktail (see Table S4 for a list of antibodies used) was resuspended in 5% DS-TBS-T + 5 μM EDTA and then passed through a 0.1 um centrifugal filter (Millipore). Tissues were stained with antibody cocktail overnight in a humidity chamber at 4°C. The following day, slides were washed twice with TBS-T, followed by PBS and then antibodies were fixed to tissue by with incubating with 2% glutaldehyde (Electron Microscope Sciences)-PBS for 5 min and neutralized with 3 volumes of 100 mM Tris pH 8.0. Slides were washed with ddH2O (2x), 70% ethanol (1x), 80% ethanol (1x), 95% ethanol (2x), and 100% ethanol (2x), air dried for 10 min and under vacuum until MIBI scanning. Multiplexed ion beam imaging data acquisition: Imaging was performed using a MIBI-TOF instrument (IonPath) with a Hyperion ion source. Xe^+^ primary ions were used to sequentially sputter pixels for a given field of view. The following imaging parameters were used: acquisition setting: 80 kHz; field size: 400 x 400 mm, 1024 x 1024 pixels; dwell time: 1 ms; median gun current on tissue: 10.5 nA Xe^+^. After image acquisition, single-channel TIFFs were extracted from raw bin files via the Angelo Lab’s toffy pipeline (https://github.com/angelolab/toffy/tree/main). Using this pipeline for all subsequent processing steps, single-channel TIFF images were mass compensated and normalized to reduce signal interference and retain comparable signal across collected FOVs. Cleaned images were visualized in ImageJ.

### Immunofluorescent staining

Kidneys were dissected and immediately placed in cold PBS. The renal capsules were removed, and the kidneys were bisected longitudinally. Tissue was briefly rinsed in Tissue-Tek OCT compound to remove excess PBS and embedded in OCT within labeled plastic cassettes, ensuring the absence of air bubbles and orienting the cut surface downward. Rapid freezing was performed by immersing the cassette in an ethanol-dry-ice slurry. Frozen samples were transferred to dry ice for temporary storage and then stored at −80 °C until sectioning. Cryosections were allowed to equilibrate to room temperature and air-dried for 1 h after removal from -80 °C if previously fixed. Tissue sections were circled using a hydrophobic barrier pen and rehydrated in PBS for 15 min. Sections were blocked for 1hr at room temperature in PBS containing 2% bovine serum albumin (BSA) and 0.1% Tween-20, with optional nuclear counterstain (e.g., DAPI). Primary staining was performed for 1 hr at room temperature using FITC-conjugated antibodies diluted in PBS containing 0.5% BSA and 0.1% Tween-20: anti-mouse IgG (1:100; Jackson ImmunoResearch, #115-096-008), anti-mouse IgM (1:300; Jackson ImmunoResearch, #115-096-075), and anti-C3 (1:75; Cappel, #55510). Sections were washed three times for 5 min each in PBS containing 0.5% BSA and 0.1% Tween-20 with gentle agitation. Slides were mounted using ProLong™ SlowFade™ mounting medium (Invitrogen, S36936) and sealed with nail polish.

### 10x single-cell RNA sequencing and analysis

Islets were purified following standard collagenase protocols as described previously (Tang et al., 2004) and dissociated by incubation in enzyme-free cell dissociation buffer (Gibco). The single-cell suspensions were stained with Live/Dead Fixable Blue (ThermoFisher) and CD45.1 eFluor506 (A20, Invitrogen) and sorted on a FACSAria II (Becton Dickinson) cell sorter for CD45^+^ cells and sent to the Genomics CoLab at the University of California, San Francisco, for single-cell library preparation and sequencing. Reads were processed and aligned using the 10x Genomics Cell Ranger count pipeline. Seurat objects for each sample were created and merged. Barcodes with a doublet score >0.15 and mitochondrial reads >5% or <200 features were removed. Preprocessing, clustering, and dimensionality reduction were performed in Rstudio (Posit team, 2023, http://www.posit.co/) using Seurat (Stuart et al., 2019). A small number of additional contaminants (doublets and non-CD45^+^ cells) were identified and removed. The object was reclustered and reprocessed. Differential genes were determined with Seurat’s “FindAllMarkers” and “FindMarkers” functions and represented as heatmaps (plotted with “pheatmap”; Kolde R, 2019, https://CRAN.R-project.org/package=pheatmap) or volcano plots (significance test.use = wilcox, plotted with “ggplot2”; Whickham et al., 2016, https://ggplot2.tidyverse.org). Cell-cell communication analysis was performed using the CellChat R package (Jin et al., 2024). Seurat object was used as input, and cell type annotations were applied to define sender and receiver populations. Overall communication patterns were visualized using “netVisual_circle” and “netVisual_heatmap” to display pathway-specific signaling intensities. The total number and relative strength of interactions were compared across genotypes and timepoints to identify context-specific shifts in intercellular communication networks.

### T cell proliferation assay

T cells from the transgenic BDC2.5 mice were enriched from spleens and control LNs (cervical, inguinal, axillary, and brachial) using the EasySep mouse naive T cell isolation kit (Cat#19765, Stemcell Technologies. Dendritic cells from either WT or Skap2^G153R/G153R^ mice were isolated using the EasySep mouse CD11c^+^ isolation kit (Cat#18780, Stemcell Technologies). 2.5 μL of CellTrace CFSE staining solution was diluted in 2.5 mL of warm PBS. The CFSE solution was added to the T cells, which were incubated for 20 minutes at 37°C in the dark. After washing and resuspending, T cells and DCs were incubated in a 96-well plate with different concentrations of p31 peptide (5 μg/mL, 2.5 μg/mL, 1.25 μg/mL, or 0 μg/mL) and placed in a 37°C (5% CO_2_) incubator for four days. Cells were washed and stained with a panel to assess T-cell proliferation. Data was analyzed using FlowJo (Treestar). Reagents listed in Table S4.

### T cell/dendritic cell conjugate formation

T cells from transgenic BDC2.5 mice were enriched from spleens and control LNs as described above and incubated with CFSE (Invitrogen) labeled dendritic cells from WT or SKAP2^G153R/G153R^ mice (isolation described above). Cells were incubated for 30 minutes at 37°C, washed with PBS, stained with anti-TCRβ for 30 minutes at 4°C, washed, and run on the BD LSRII flow cytometer.

### Confocal analysis

MatTek glass-bottom dishes (MatTek, P35GC-1.5-14.C) were prepared as described in reference (Kersten et al., 2023). CTV-labeled dendritic cells from WT or *Skap2* G153R cells were isolated (described above) and incubated with 5 μg of p31 peptide at 37°C for 30 minutes. The slides were washed with PBS. Right before imaging, T cells from BDC2.5 mice (resuspended in 0.1% agarose) were added to the chambers, and slides were loaded for imaging. Videos were acquired using a SORA spinning disk confocal microscope acquired at 20x magnification. Samples were kept at 37°C with CO_2_. Image analysis was performed using the Imaris (Bitplane) software.

### Scratch assay

Bone marrow from WT and *Skap2* G153R mice was isolated, and the single-cell suspensions were layered onto a 62% Percoll gradient and spun at 500g for 30 min with the brakes off. Mononuclear cells were collected from the interface of the gradient, washed, and plated on Valmark plates in macrophage media (αMEM containing 10% FCS, Pen-strep, and 100 ng/ml M-CSF (supernatant harvested from a 3T3 fibroblast cell line engineered to overexpress M-CSF) for 48 hours. After 48 hours, nonadherent cells were harvested and plated in 6 well tissue culture plates at 5 X 10^5^ cells per plate and rested at 37°C overnight. The next day, confluent plates were scratched with a P200 pipette tip, and time points were taken at 0, 3, 6, 9, 12, and 24 hours after the scratch. Plates were fixed in 4% PFA in PBS with DAPI and imaged on the EVOS cell imaging system.

### Bead-based stimulation of integrin α_v_ in macrophages

Bead preparation and binding assay were performed and imaged as described previously.(Alenghat et al., 2012) Following 20 min of bead stimulation, cells were stained with 1:500 Alexa Fluor™ 546 Phalloidin (Cat#A22283, ThermoFisher) and primary antibody anti-SKAP2 and 1:1000 of secondary antibody Goat-Anti-Rabbit IgG Alexa Fluor™ 488 (Cat#A-11008, ThermoFisher), then directly imaged on a Leica SP5 confocal microscope.

### ROS assay

Isoluminol-enhanced chemiluminescence to measure ROS as described previously (Abram et al., 2014). Briefly, bone marrow-derived neutrophils purified using a 62% Percoll gradient, plated on a poly-RGD coated 96-well plate, and stimulated with or without TNFα were read by a chemiluminescence plate reader to determine ROS production (Molecular Devices Spectramax M5). Ibrutinib (Fisher Scientific, 936563-96-1) or DMSO (control) was added at different concentrations and incubated at 37°C for 15 minutes, then washed, and the stimulus was added (see ROS assay methods).

### ICAM binding assay

Bone marrow (BM) cells were isolated from WT and KI mice. BM cells were incubated for 10 min at room temperature (RT) with 20 µg/mL ICAM-1/Fc (R&D systems, cat. no AVI-7201-05) and APC-conjugated anti-human IgG1 (Fc specific, 9042-11, Southern Biotechnology). Without washing out, cells were left untreated or stimulated with leukotriene B4 (LTB4, 150 ng/mL) or phorbol myristate acetate (PMA, 100nM) for 15 min at 37°C. After stimulation, cells were washed, stained with Ly6G–PE-Cy7 (1:200) and Zombie UV (1:200), and analyzed by flow cytometry.

### Pristane-induced autoimmunity

Pristane (Sigma P9622 0.5 ml) was administered intraperitoneally as a single dose. Peripheral blood was collected monthly post-injection in gel CAT 1.1 tubes (Sarstedt) and allowed to clot at RT for ∼45min at room temperature. Tubes were centrifuged for 2 mins at 6,000xg to obtain serum. Sera were diluted 1:40 and applied to Kallestad HEp-2 slides (Bio-Rad). Anti-nuclear antibodies were detected using FITC-conjugated goat anti–mouse IgG (Jackson ImmunoResearch Laboratories). For ANA quantification, the mean pixel intensity of cells in the green channel was calculated from a DAPI counterstain-defined mask using ImageJ/Fiji.

### Statistical methods

All statistical analysis was performed using Prism 10 (GraphPad Software; www.graphpad.com). Figures display mean ± SE values. All data were compared using two-way Anova or T-testing, with the exception of diabetes onset curves, which were analyzed using Log-rank testing (Mantel-Cox test). A p-value <0.05 was considered statistically significant. The number of replicates for each experiment is indicated in the figure legends.

## Acknowledgments

This work was supported by the Leona M. and Harry B. Helmsley Charitable Trust (G-2018PG-T1D018, G-2003-04376, G-2404-06823), the National Institute of Diabetes and Digestive and Kidney Diseases (R01-DK-104942, P30-DK-020595, T32DK007418), the National Institute of Allergy and Infectious Diseases of the National Institutes of Health (R01 AI170841), and a private donor.

We acknowledge the Parnassus Flow Cytometry CoLab PFCC (RRID:SCR_018206) supported in part by Grant NIH P30 DK063720 and by the NIH S10 <u>1S10OD021822-01 (Aurora Cytek),</u> NIH S10 <u>1S10OD021822-01</u> (Aria sorters and analyzers) and NIH S10 <u>1S10OD025187-01</u> (MIBIscope). We would like to acknowledge the Biological Imaging Development CoLab (BIDC) staff at UCSF Parnassus Heights for their training and support in using the Nikon CSU-W1 SoRa Spinning Disk Microscope, supported by the grant S10OD028611-01. Additionally, Austin Edwards has been exceptionally helpful in providing training and assistance with data analysis. We thank Yongmei Hu for assistance with the maintenance of the multiple mouse strains used in this study. We thank Alyssa Indart, UCSF, for help with BDC2.5 transgenic mice. Illustrations were created using Biorender.com. The authors thank the family for participation in this research study (https://precisiont1d.uchicago.edu/)

## Author Contributions

C.M.T., C.E.C., C.L.A., S.P., and J.B. conducted the majority of experiments within the manuscript. J.L.M. generated the SKAP2^G153R/G153R^ mouse model. N.R., L.S., and W.D. conducted initial experiments on the mouse model and on cell lines containing the *Skap2* mutation. W.T. provided computational support for scRNA-seq analysis and phenotype confirmation. L.R.L and L.H.P. are care providers for the index *SKAP2* patient. I.P., M.S.G., M.S.A., and C.A.L. oversaw the work, did the writing and editing of the manuscript, and shared the grant funding for this project.

## Notes

### Competing Interest Statement

The authors have declared no competing interest.

