## Supplemental Materials for "Gain-of-function mutation in *SKAP2* leads to type 1 diabetes and broader autoimmunity through hyperactive integrin signaling in myeloid cells"

### Supplemental Data

#### FIGURE S1

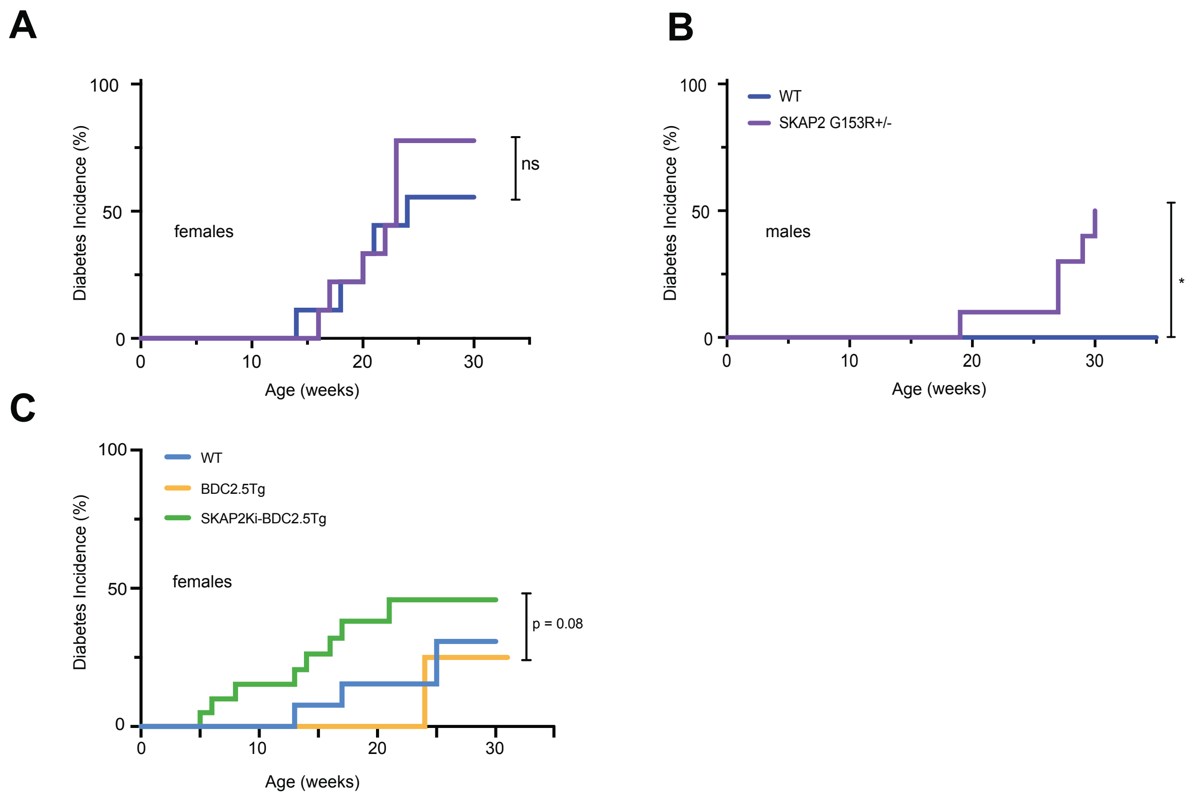


Supplemental Figure 1. **The SKAP2 G153R mutation accelerates autoimmune diabetes and breaks tolerance in the BDC2.5 transgenic model.**

**(A, B)** Cohorts of WT and SKAP2$^{G153R/+}$ female **(A)** and male **(B)** mice were monitored for blood glucose levels weekly. Diabetes incidence (blood glucose > 200 mg/dl on two tests separated by one week) incidence is shown onset and incidence over 30 – 40 weeks is shown (females: WT, n=9; SKAP2$^{G153R/+}$, n=10; males: WT, n=10; SKAP2$^{G153R/+}$, n=10). **(C)** Cohorts of female WT, BDC2.5 transgenic mice (on the NOD genetic background) and SKAP2$^{G153/+}$ - BDC2.5 transgenic mice were monitored for blood glucose levels weekly as above. Cohort sizes: WT, n=15; BDC2.5 Tg, n=10; SKAP2$^{G153R/+}$- BDC2.5, n=20). Diabetes onset curves were analyzed using Log-rank testing (Mantel-Cox test); *=$p<0.05$

### FIGURE S2

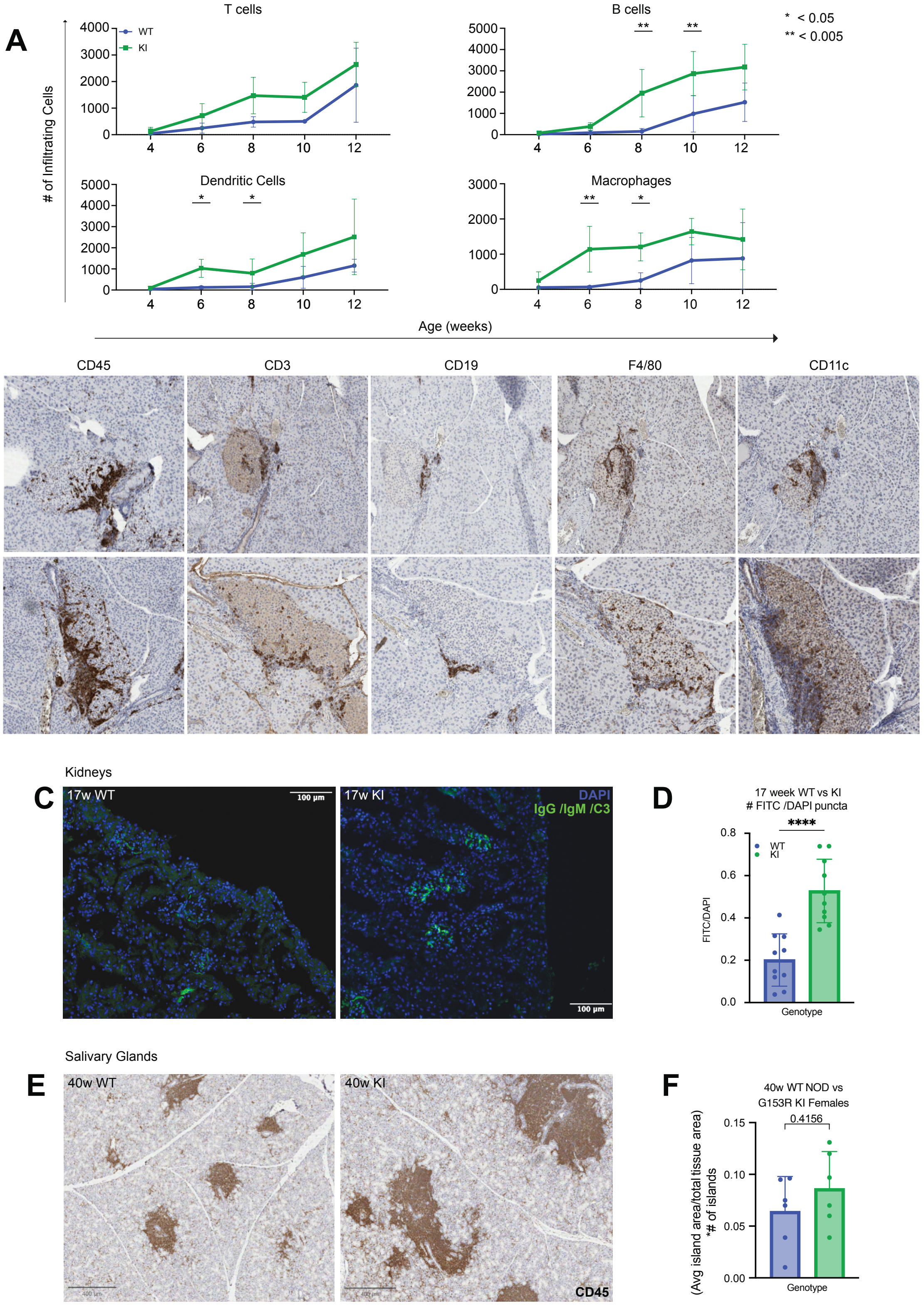
A
T cells
WT
KI
B cells
* < 0.05
** < 0.005
### of Infiltrating Cells
Dendritic Cells
Macrophages
Age (weeks)
B
CD45
CD3
CD19
F4/80
CD11c
6w WT
6w KI
Kidneys
C
17w WT
17w KI
DAPI
IgG /IgM /C3
100 µm
D
17 week WT vs KI
### FITC /DAPI puncta
****
FITC/DAPI
Genotype
Salivary Glands
E
40w WT
40w KI
CD45
F
40w WT NOD vs
G153R KI Females
0.4156
(Avg island area/total tissue area)
*# of islands
Genotype

Supplemental Figure 2. **Histologic analysis of NOD WT vs SKAP2 KI mice.**

**(A)** Pancreases from female WT and SKAP2$^{G153R/G153R}$ (KI) mice were prepared for histology as described in the methods – staining for CD45, CD3, CD19, F4/80, and CD11c (see Table S4 for antibodies used) to detect total leukocytes, T cells, B cells, macrophages, and dendritic cells, respectively. Pancreases from mice of the indicated age (4, 6, 8, 10, 12) were used. Scanned images of 10 different islets were analyzed for leukocyte infiltration by QuPath software to quantify cell numbers. Each age cohort had n=5 mice each. Data for each cell type at each time point were averaged and plotted as individual line graphs. P values were calculated by two-way ANOVA. *=p<0.05; **=p<0.01. **(B)** Representative images of pancreatic islets, stained for the above markers, from 6-week-old female WT and KI mice. **(C–D)** Immune complex deposition in the kidneys of 17-week-old WT and KI mice was assessed by IgG/IgM/C3 immunofluorescence staining and quantification. **(E–F)** $CD45^+$ immune infiltration in salivary glands of 40-week-old WT and KI mice and corresponding quantification. In bar graphs, each dot represents samples from a single mouse. P values were calculated by unpaired t-tests. Data represent the difference between means ± SEM; ****=p<0.0001.

### FIGURE S3

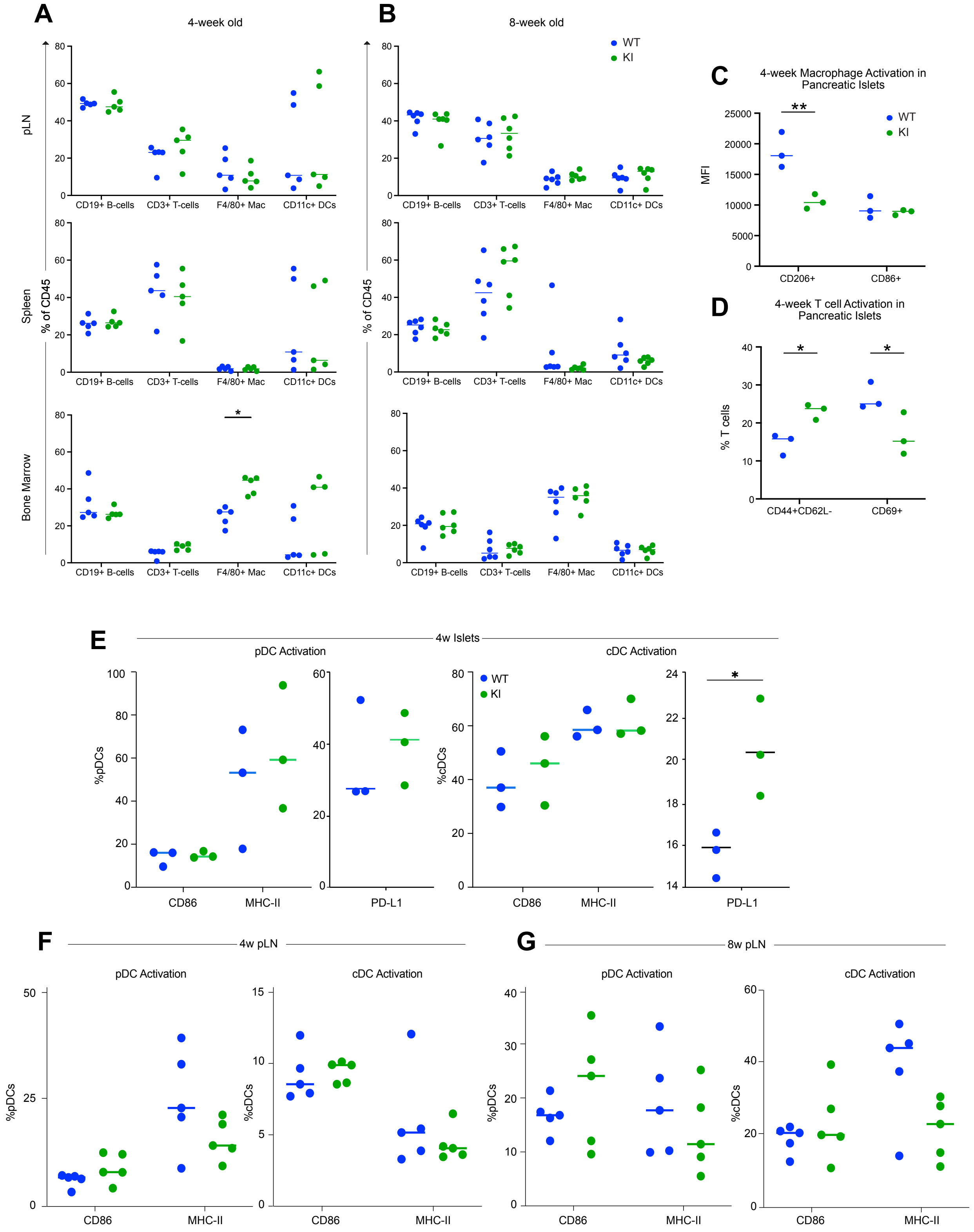
A
4-week old
pLN
Spleen
Bone Marrow
% of CD45
CD19+ B-cells
CD3+ T-cells
F4/80+ Mac
CD11c+ DCs
B
8-week old
WT
KI
C
4-week Macrophage Activation in Pancreatic Islets
MFI
CD206+
CD86+
D
4-week T cell Activation in Pancreatic Islets
% T cells
CD44+CD62L-
CD69+
E
4w Islets
pDC Activation
cDC Activation
%pDCs
%cDCs
CD86
MHC-II
PD-L1
F
4w pLN
G
8w pLN

Supplemental Figure 3. **Flow cytometric analysis of WT vs SKAP2 KI mice.**

Multiparameter flow cytometry was conducted on pancreatic draining lymph node (pLN), spleen, and bone marrow tissues isolated from four or eight-week-old female WT versus SKAP2$^{G153R/G153R}$ (KI) mice. Cell suspensions from the tissues were stained with antibodies listed in Table S4, then analyzed on an Aurora Spectral Flow cytometer. **(A, B)** The percentages of B cells, T cells, macrophages, and dendritic cells within each tissue are shown (n=5 mice per group – each dot represents one animal). **(C)** The mean fluorescence intensity (MFI) of the marker CD206 or CD86 on F4/80$^+$ macrophages from isolated pancreatic islets of four-week-old mice is shown on the upper panel. **(D)** The percentages of activated (CD44$^+$CD62L$^-$) or replicating (CD69$^+$) T cells from the same islet samples are shown. **(E)** Markers of DC activation (MHC II, CD86, and PD-L1) were stained on islet cell preparations, and the percentages of CD11c$^+$ pDCs or cDCs displaying these markers are shown in samples from four-week-old NOD WT and SKAP2$^{G153R/G153R}$ (KI) mice. **(F)** Markers of DC activation in cDCs and pDCs from pancreatic draining lymph nodes (LN) are shown. Cells from Islets and draining LN were pooled from n=3 mice per group for cell suspension generation. Note that enumerating the total number of immune cells within the isolated islets by flow cytometry was not useful, as many immune cells remained within the pancreatic tissue after islet removal. Histology proved more reliable for total cell counting. P values were calculated using two-way ANOVA analysis; *=p<0.05; **=p<0.01.

### FIGURE S4

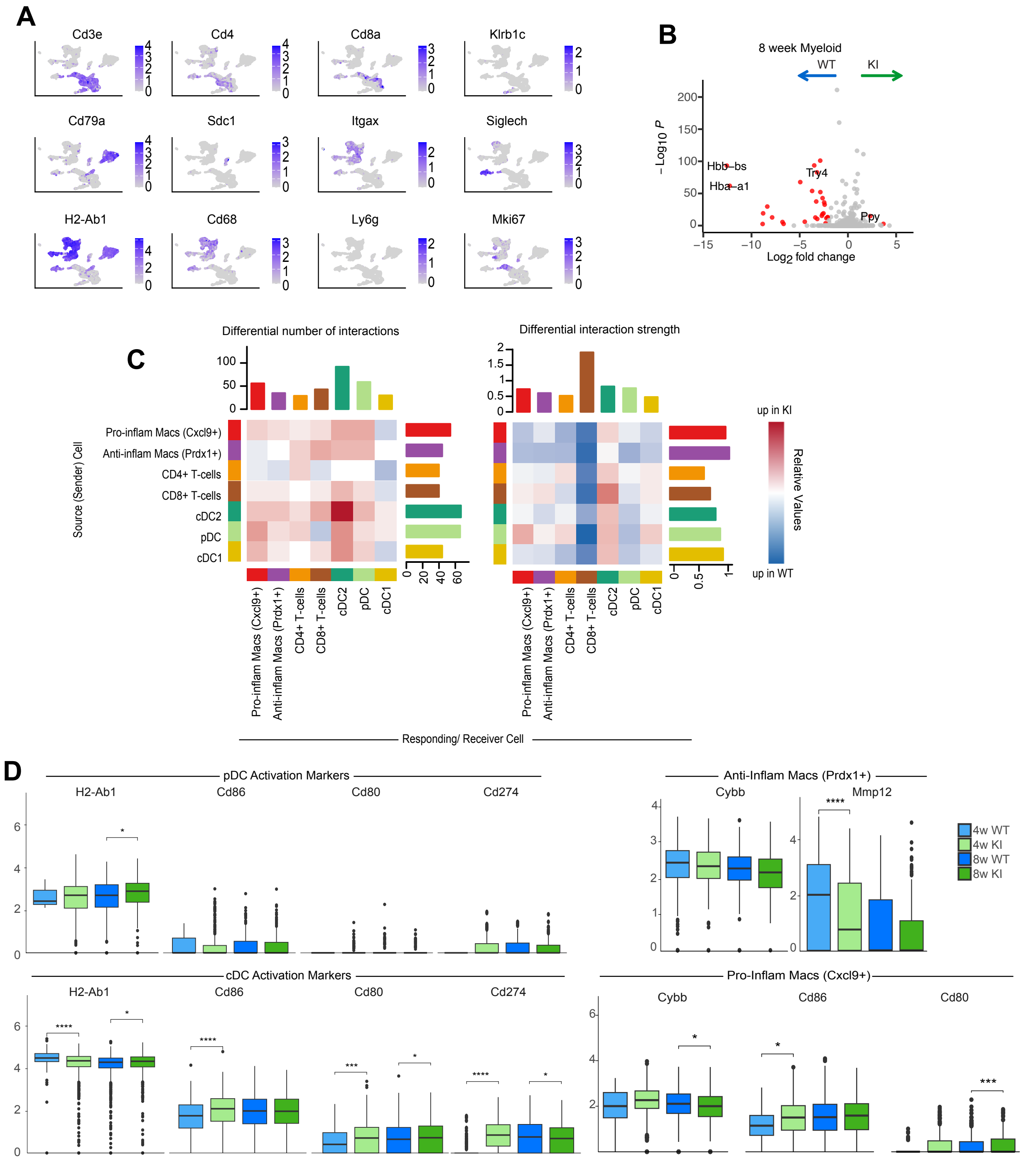


Supplemental Figure 4. **Gene set, Cell Chat, and individual transcript analysis of immune cells from islets of WT versus SKAP2$^{G153R/G153R}$ KI mice.**

**(A**) Individual feature plots for specific lineage-defining transcripts (shown on the overall plot of islet cells from four-week-old SKAP2$^{G153R/G153R}$ mice). (**B)** Gene set analysis (GSEA) of all CD45$^{+}$ cells from eight-week-old WT versus SKAP2$^{G153R/G153R}$ (KI) mice, with differences plotted in a volcano plot. Red dots indicate genes showing $\log_2$ fold difference > 0.3 and $-\log_{10}$ > 1.3 (giving a $p<0.05$), which are default cutoffs in Seurat/Volcano packages. This analysis indicates that by eight weeks of age, few differences in leukocyte gene expression patterns were observed between WT and KI mice. **(C)** Cell Chat data plotted as a box plot, showing the number of predicted cellular interactions and interaction strength between islet immune cells in WT and SKAP2$^{G153R/G153R}$ (KI) mice. Differential interactions are represented by red versus blue boxes in the two plots. The relative increase in red boxes on the left suggests that the KI immune cells will form more interactions, while the blue boxes on the right indicate that the WT cells will exhibit stronger interactions. **(D)** Transcript read counts for various activation makers and inflammatory proteins in sorted DC and macrophage subsets. Shown are counts for the transcripts for encoding MHC II, CD80, CD86, and CD274 in DC subsets (cDCs versus pDCs), along with *Cybb* (encoding NOX2) and Mmp12 in macrophage subsets. Each dot represents the read count for the indicated gene in a single cell. P values were calculated using two-way ANOVA analysis; *=p<0.05; **=p<0.01, ***=p<0.001, ****=p<0.0001.

### FIGURE S5

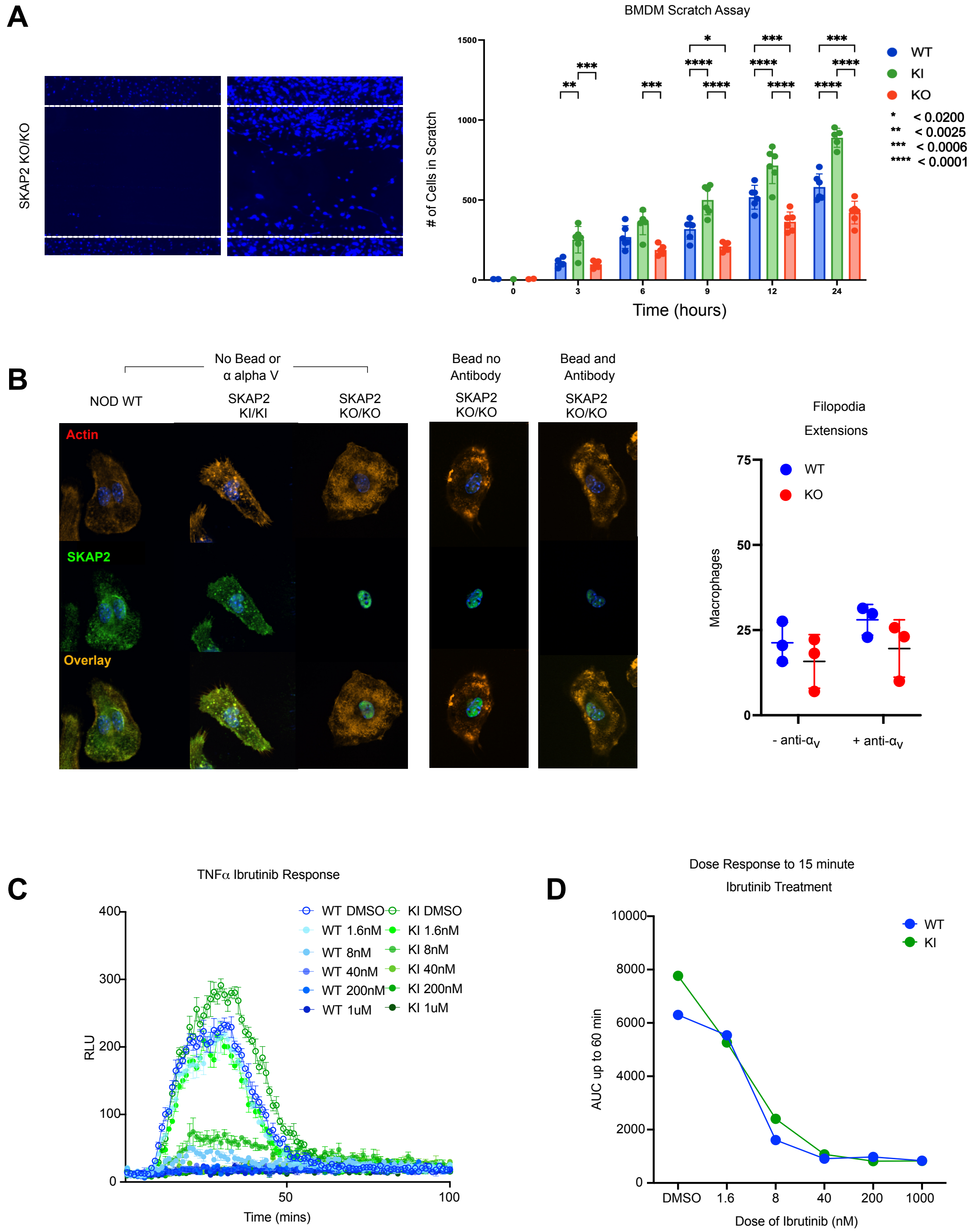


Supplemental Figure 5. **Comparison of integrin signaling in WT versus SKAP2 G153R versus *Skap2*$^{-/-}$ macrophages and neutrophils.**

**(A)** Bone marrow-derived macrophages (BMDM) from female WT, homozygous *Skap2* G153R (KI), and S*kap2*$^{-/-}$ (KO) mice were expanded on tissue culture plates, and scratch assays were performed as described in the methods. The left panel shows migration responses in *Skap2*$^{-/-}$ cells, compared with the micrographs in Fig. 6a. The right panel shows a time course of WT, KI, and KO cell migration into scratched areas. Each dot represents a separate scratch culture. Cell counting was performed using QuPath software. The statistical significance of differences between groups is shown. **(B)** Confocal micrographs of WT, KI, or KO BMDMs plated on glass coverslips, then stimulated with uncoated polystyrene beads or with beads coated with anti-$\alpha$v integrin antibody for 20 min, then fixed and stained for F-actin with phalloidin (red) and anti-Skap2 (green). Merged images show the overlap of the stains in yellow. Filopodia extensions were quantified in unstimulated versus stimulated WT and KO cells. **(C)** Bone marrow-derived neutrophils from WT or KI mice were pre-treated with the indicated doses of ibrutinib for 15 min, then plated on poly-RGD-coated wells with TNFα. ROS production was monitored by isoluminol reduction as described in the methods. **(D)** The total area under each curve (from 0 to 60 min) in panel c is shown for each ibrutinib dose. These data indicate that WT and KI neutrophils show the same dose-response to ibrutinib inhibition of integrin signaling in this ROS production assay.

#### FIGURE S6

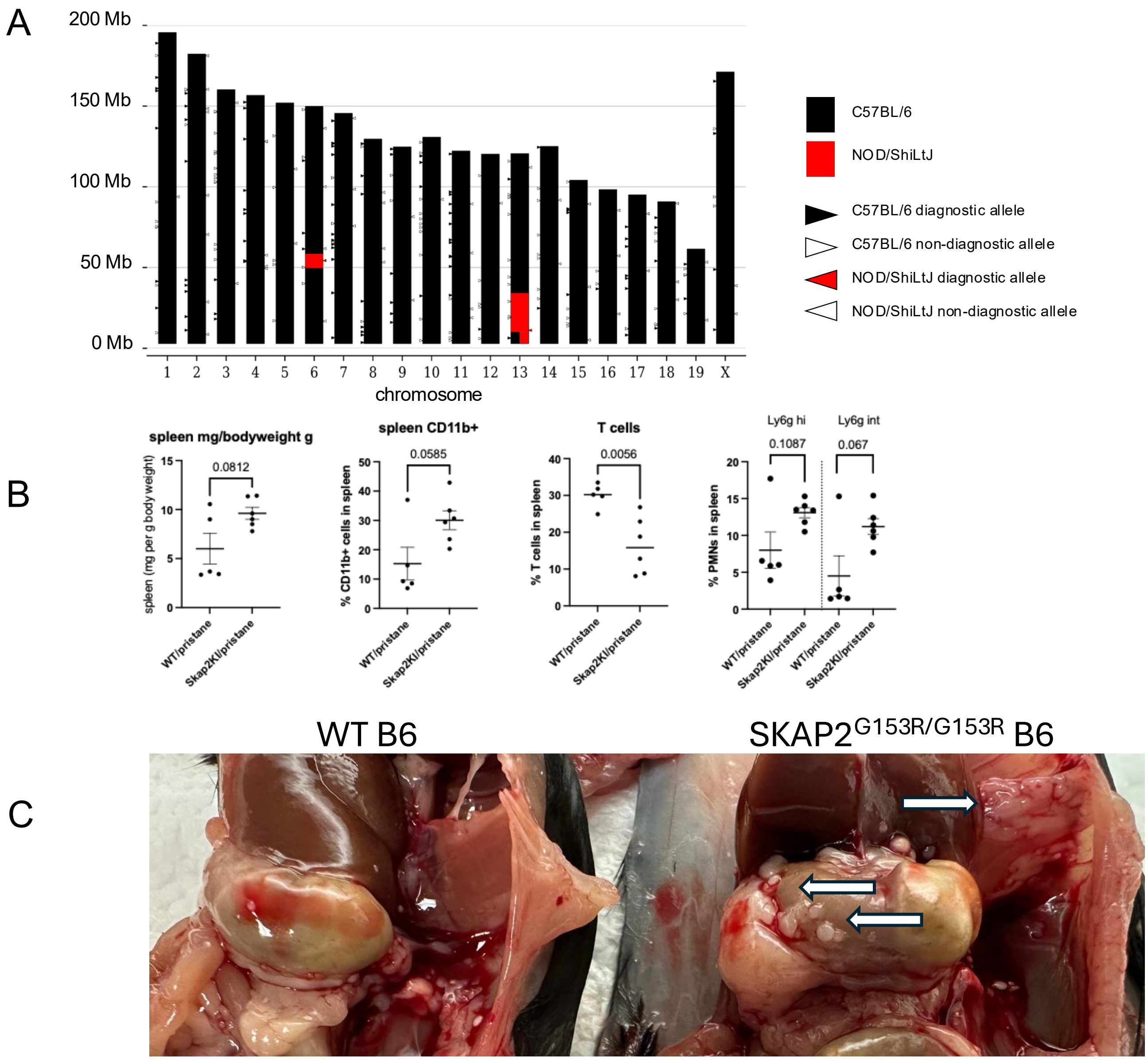


Supplemental Figure 6. **Pristane induced inflammation in WT B6 versus SKAP2$^{G153R/G153R}$ B6 mice.**

**(A)** SKAP2$^{G153R/G153R}$ NOD mice were backcrossed to C57BL/6 for 12 generations, with the mutation confirmed by genotyping. The genetic background was analyzed by SNP genotyping of tail genomic DNA from breeders. Six breeders were tested and found to have an average of 5%

NOD and 95% C57BL/6 background strains. Summary data from one of the breeders is shown above. Mouse *Skap2* is found on chromosome 6 at ~51.8Mbp, and all mice showed NOD diagnostic markers in this region. Other regions of NOD were found on chromosome 8 (3/6 mice) and 13 (6/6 mice). **(B)** Spleen weights and cell counts in WT B6 versus SKAP2$^{G153R/G153R}$ mice at three months following pristane injection. P values were calculated using two-way ANOVA analysis. **(C)** Photomicrograph of peritoneal lymphoid aggregates (shown by white arrows) in WT B6 versus SKAP2$^{G153R/G153R}$ mice at three months following pristane injection.

**Supplemental Table 1:** List of antigens on array heat maps, from top to bottom, on Fig 2C:

| | | |
|---|---|---|
| AChR3 | ErbB3&B4 | RSVP antigen |
| AFP | ERP29 | Rubella virus grade III antigen |
| ALDOC | Ferritin | Rubella virus grade IV antigen |
| Aldolase (muscle) | Flagellin CBir1 | Rubeola antigen |
| Alpha-2-macroglobulin-like 1 | FUCA1 | Saccharomyces Cerevisiae |
| alpha-fodrin | Galactocerebroside | Salivary Gland Protein 1 (SP1) |
| Angiotensin II type 1 Receptor | GBM | SCCA |
| ANXA1 | Glutamate decarboxylase-65/GAD2 | SOX2 |
| Apolipoprotein B/E | HEPATITIS A ANTIGEN | Sphingomyelin |
| Azurocidin | HPV E6 (16) | Tetanus toxoid |
| BAFF | HPV E7 (16) | TGF-beta1 |
| beta-glucuronidase | HPV E7 (18) | Thrombopoietin (TPO) |
| BRCA1 | HSV-1 ANTIGEN | Thyroglobulin |
| BRCA2 | HSV-2 ANTIGEN | Tissue Transglutaminase (TTG) |
| C-MYC | IFN-alpha2 | TNF-alpha |
| CA15-3 | IFN-beta | TNF-beta |
| CA19-9 | IFN-gamma | Toxoplasma |
| CA27-29 | IL-1alpha | Troponin I |
| CA50 | IL-1beta | Troponin I-T-C ternary complex mixture |
| CA72-4 | IL-2 | TWEAK (CD255) |
| CA125 | IL-6 | uPA |
| CA242 | IL-8 | VEGF-165 |
| CAGE | IL-12 (p70) | VZV antigen |
| Calmodulin (brain) | IL-17A | VZV grade II antigen |
| Calreticulin(rabit calreticulin) | lactoferrin | |
| Carbonic Anhydrase 6 (CA6) | LC1 | |
| Catalase | Lysosomal Associated Protein 2 (LAMP-2) | |
| Cathepsin G | MAG | |
| CCL2 | MAGEA3 | |
| CCL3 | MBP | |
| CCL5 | MMP-2 | |
| CCL11 | NSE | |
| CDK2 | Nucleosome | |
| CEA | NY-ESO-1 | |
| Cholinergic Receptor Muscarinic 3 (CHRM3) | OmpC | |
| CMV-G | Ox40L | |
| CMV-M | P53 | |
| CMV EXT | Parotid Secretory Protein (PSP) | |
| CMV GRADE III ANTIGEN | PD1 | |
| Complement Factor Hq | PD-L1 | |
| CTLA4 | Peptidyl Arginine Deiminases 1&2&3&4 | |
| CXCL10 | Phosphatidyl-l-serine | |
| Cytokeratin 19 Ag | PKM2 | |
| dsDNA | Prostate Specific Antigen (PSA) | |
| EBV EBNA1 | Prostate Specific Membrane Antigen (PSMA) | |
| EGF Receptor | Prostatic Acid Phosphatase | |
| Elastase | PSMA | |
| Endothelial Cell Extract | rhHSPG2 | |
| Enolase | Rotavirus SA-11 | |
| ErbB2 | RSV antigen | |

**Supplemental Table 2:** List of genes on PhIP-seq heat maps, from top to bottom, on Fig 2F:

| |
|---|
| 4931409K22Rik |
| Actr10 |
| Apc |
| Blnk |
| Bnc1 |
| Cdc37 |
| Cftr |
| Chd4 |
| Cherp |
| Col11a2 |
| Cp |
| D6Ertd527e |
| Dlx5 |
| Fbxo28 |
| Fhad1 |
| Fhod3 |
| Gm33989 |
| Kirrel |
| Kmt2d |
| Lhx3 |
| Mast2 |
| Mpdz |
| Mre11a |
| Myrf |
| Nrg2 |
| Olfr1129 |
| Osbpl6 |
| Ppfibp2 |
| Prkab1 |
| Psmb11 |
| Ralgapa1 |
| Reep6 |
| Serpina3f |
| Shank3 |
| Slf2 |
| Snrpb |
| Snrpn |
| Spag8 |
| Speg |
| Tex22 |
| Trerf1 |
| Uba1 |
| Vldlr |
| Zfp207 |
| Zfp78 |
| Zfp846 |
| Zmym4 |

**Supplemental Table 3:** List of genes on PhIP-seq heat maps, from top to bottom, on Fig. 7A:

| | |
|---|---|
| 4930452B06Rik | Tenm4 |
| Abca4 | Tet3 |
| Arid1b | Trip10 |
| Asxl3 | Ttc23 |
| Atad2b | Vwc2l |
| Atp10b | Vwc2l |
| Atp2b3 | Wnt7b |
| Bpifb4 | Zcchc2 |
| C1qa | |
| Carmil1 | |
| Cbarp | |
| Cntn5 | |
| Crk | |
| Ddi2 | |
| Eppk1 | |
| Fam120c | |
| Fnbp1 | |
| Gabra1 | |
| Gm46637 | |
| Golga4 | |
| Grhl2 | |
| Grhl3 | |
| Hnrnpa1l2-ps2 | |
| Hormad2 | |
| Ikbip | |
| Kcnb1 | |
| Lrrfip1 | |
| Man1a | |
| Mcc | |
| Mrap2 | |
| Mre11a | |
| Nhs | |
| Nom1 | |
| Numa1 | |
| Pdgfb | |
| Plch2 | |
| Plec | |
| Pop1 | |
| Ppan | |
| Prex1 | |
| Ptges3 | |
| Rad51b | |
| Rictor | |
| Rptor | |
| Rtf1 | |
| Setd2 | |
| Sned1 | |
| Srcin1 | |
| Tenm4 | |
| Tet3 | |
| Trip10 | |
| Ttc23 | |
| Vwc2l | |
| Wnt7b | |
| Zcchc2 | |
| Srcin1 | |

**Supplemental Table 4:** Antibody staining reagents used:

#### Immunohistochemistry antibodies

| Marker | Clone | Vendor | Dilution from stock |
|---|---|---|---|
| CD45 | EPR20033 | Abcam | 1:1000 |
| CD3 | EPR22667-12 | Abcam | 1:500 |
| CD19 | EPR23174-145 | Abcam | 1:500 |
| F4/80 | D2S9R | Cell Signaling | 1:300 |
| CD11c | D1V9Y | Cell Signaling | 1:300 |

#### MIBI reagents

| Isotope | Metal | Antibody | Vendor | Clone | Micrograms/$10^6$ cells |
|---|---|---|---|---|---|
| 31 | P | Phosphorous (P) | | | 1 |
| 39 | K | K | | | 1 |
| 40 | Ca | Ca | | | 1 |
| 56 | Fe | Iron (Fe) | | | 1 |
| 89 | Y | Histone H3 | CST | DIH2 | 0.25 |
| 141 | Pr | FoxP3 | Abcam | Polyclonal | 5 |
| 143 | Nd | CD4 | Abcam | EPR19514 | 1.2 |
| 144 | Nd | Cd11c | CST | D1V9Y | 1.2 |
| 150 | Nd | Granzyme B | Abcam | D6E9W | 1.2 |
| 152 | Sm | SKAP2 (biotin) | Invitrogen | Polyclonal | 1.2 |
| 153 | Eu | Ki67 | Abcam | SP6 | 1.2 |
| 155 | Gd | CD11b | abcam | EPR1344 | 1.2 |
| 156 | Gd | F4/80 | ab254201 | D2S9R | 1.2 |
| 157 | Gd | CD103 | Ms | EPR22590 | 1.5 |
| 158 | Gd | CD8 | Ms | D4W2Z | 1 |
| 159 | Tb | CD3e | Abcam | EPR22667-12 | 1 |
| 162 | Dy | Glucagon | CST | EPR3070 | 1 |
| 163 | Dy | Vimentin | CST | D21H3 | 0.75 |
| 165 | Ho | Pax5 | Abcam | D7H5X | 1.5 |
| 166 | Er | Alpha-SMA | Abcam | SP171 | 1 |
| 167 | Er | CD19 | Abcam | EPR23174 | 2 |
| 172 | Yb | Trypsin | Abcam | EPR19498-43 | 2 |
| 173 | Yb | SIRP$\alpha$ | CST | EPR22590 | 2 |
| 174 | Yb | Beta-Tubulin | Abcam | D3U1W | 3 |
| 175 | Lu | CD45 | Abcam | EPR20033 | 1 |
| 176 | Lu | Insulin | Abcam | EPR173591 | 1 |

#### Flow markers and fluorophores

| Marker | Fluorophore | Clone | Vendor | μg/10$^6$ cells |
|---|---|---|---|---|
| CD44 | BUV661 | IM7 | BD Biosciences | 0.125 |
| CD4 | BUV805 | GK1.5 | BD Biosciences | 0.125 |
| CD117 | PE-Cy5 | 2B8 | BioLegend | 0.25 |
| F4/80 | BUV563 | T45-2342 | BD Biosciences | 0.25 |
| CD64 | Brilliant Violet 605 | X54-5/7.1 | BioLegend | 0.25 |
| CD206 | Brilliant Violet 711 | C068C2 | BioLegend | 0.25 |
| CD103 | AF488 | 2.00E+07 | BioLegend | 0.25 |
| TCRgd | PerCP-eFluor710 | GL3 | Invitrogen | 0.25 |
| CX3CR1 | PE | SA011F11 | BioLegend | 0.25 |
| Siglec-F | PE-CF594 | E50-2440 | BD Biosciences | 0.25 |
| CD19 | BUV395 | 1D3 | BD Biosciences | 0.5 |
| CCR2 (CD192) | BUV496 | 475301 | BD Biosciences | 0.5 |
| CD69 | BUV737 | H1.2F3 | BD Biosciences | 0.5 |
| Ly-6C | SuperBright 436 | HK1.4 | Invitrogen | 0.5 |
| CD62L | Brilliant Violet 570 | MEL-14 | BioLegend | 0.5 |
| CXCR2 (CD182) | Alexa Fluor 647 | SA044G4 | BioLegend | 0.5 |
| CD8 | Alexa Fluor680 | Polyclonal | Bioss | 0.5 |
| CD11b | APC-Cy7 | M1/70 | Tonbo Biosciences | 0.5 |
| MHC II I-Ak | BUV615 | 10-3.6 | BD Biosciences | 1 |
| B220 | Pacific Blue | RA3-6B2 | BD Biosciences | 1 |
| CD11c | BV480 | N418 | BD Biosciences | 1 |
| CD45.1 | eFluor506 | A20 | Invitrogen | 1 |
| CD86 | BV650 | GL-1 | BioLegend | 1 |
| PD-L1 | AF532 | MIH5 | Novus | 1 |
| CD66a | PerCP-Cy5.5 | Mab-CC1 | BioLegend | 1 |
| Ly-6G | AF594 | 1A8 | BioLegend | 1 |
| NKp46 | PE-Cy7 | 29A1.4 | Invitrogen | 1 |
| CD3 epsilon | PerCP | 145-2C11 | BD Biosciences | 1.5 |
| CD16/CD32 | None | 93 | BioLegend | 0.5 |
| Live/Dead Fixable Blue | | | ThermoFisher | 5 |

#### T-cell proliferation reagents

| Marker | Fluorophore | Clone | Vendor | Concentration |
|---|---|---|---|---|
| Live/Dead FB | | | Invitrogen | 5 |
| CFSE | | | Invitrogen | 1 |
| CD4 | APC-Cy7 | GK1.5 | Fisher Scientific | 0.5 |
| Thy1.1 | APC | OX7 | Fisher Scientific | 0.5 |
